# Tumor cell dissemination is facilitated through regulatory T cell-driven extracellular matrix remodeling

**DOI:** 10.1101/2025.10.25.684515

**Authors:** Ailen D. Garcia-Santillan, Jessanne Y. Lichtenberg, He Shen, Jasmine M. Rodriguez, Sandra Makar, Jonathan Barra, Nicholas M. Clark, Wei Du, Leandro M. Martinez, Ashley D. Hadjis, Taylor Calicchia, Mikhail G. Dozmorov, Jose J. Bravo-Cordero, Amy L. Olex, Priscilla Y. Hwang, Paula D. Bos

## Abstract

Regulatory T (Treg) cells function to enforce peripheral tolerance and are potent suppressors of tumor immunity. In breast cancer, we have shown that Treg cells promote tumor growth by favoring alternative activation of macrophages via suppression of IFN-γ. The tumor-associated extracellular matrix (ECM) is a key regulator of metastatic dissemination and is emerging as a critical regulator of tumor immunity. However, the reciprocal effects of the immune system on the ECM remain elusive. Using a combination of murine *in vivo*, *ex vivo* and bioengineering models we describe Treg cell-dependent changes in the tumor ECM that facilitate tumor cell migration and metastatic dissemination. Importantly, Treg cell-dependent matrisome signatures correlate with delayed survival advantage in human breast cancer samples, and are upregulated in tumor-associated macrophages (TAMs). Further, both IFN-γ and IFN-γ-sensitivity in TAMs contribute to structural and functional changes of the ECM. This work underscores a previously unrecognized role of Treg cells on the ECM that facilitates metastasis and is consistent with tissue Treg cell emergent function as critical regulators of tissue repair and cancer.

## Introduction

Breast cancer mortality has decreased by 40% in the past 3 decades, however metastatic breast cancer remains an incurable disease and takes more than 40,000 lives every year only in the United States ^1,2^. Historically perceived as an immunologically cold tumor, the success obtained through immune checkpoint blockade (ICB) in other solid tumors was not immediately mirrored in breast cancer patients. A wide range of responses have been observed when evaluating the benefit of ICB combination therapy, underscoring the heterogeneity and strong immune suppressive breast cancer microenvironment, which remains a formidable obstacle to overcome in order to harness the full power of the immune system ^3-5^.

Regulatory T (Treg) cells are a subset of CD4 T lymphocytes that express the lineage transcription factor Foxp3 and are the main enforcers of peripheral tolerance ^6^. In human breast cancer patients, their frequency is increased in tumor and peripheral blood and their presence is negatively correlated with relapse-free and overall survival ^6-9^. In murine models, we previously showed that transient ablation of Treg cells results in significant impairment of mammary tumor growth and metastasis burden ^10^. Furthermore, we found that this anti-tumor effect was dependent on interferon gamma (IFN-γ) and tumor-associated macrophages (TAMs) ^11^.

The extracellular matrix (ECM), a highly organized non-cellular network consisting of structural proteins, growth factors, cytokines and other secreted molecules, is highly dynamic and plays critical roles during tumor initiation, progression and metastasis ^10,11^. Moreover, it is now apparent that the ECM is an important immunosuppressive component of the tumor microenvironment (TME), contributing to T cell exclusion and modulating the tumor-immunity cycle through physical and molecular forces ^12-14^. Growing evidence suggests that TAMs are capable of remodeling the tumor-associated ECM, however the effects of immune cells on the ECM remain largely unexplored ^12-14^.

In this work, we sought to assess whether Treg cell-mediated immune suppression is associated with changes in the ECM, and whether those ECM changes are functionally meaningful and contribute to the strong tumor promoting effects derived from the intratumoral Treg cell population. Using spontaneous and transplantable models of breast cancer, we describe remodeling of the ECM upon Treg cell targeting, primarily of collagens, that are directly responsible for cancer cell invasive behavior and metastatic dissemination. Furthermore, Treg cell ablation-driven changes in the matrisome genes correlate with delayed survival benefit in human breast cancer samples. Bioinformatic analysis of single cell RNA sequencing profiles of primary tumors reveals this signature is prominently expressed in tumor associated macrophages and upregulated in Treg cell ablated samples. Mechanistically, antibody neutralization or genetic deletion of IFN-γ, or genetic silencing of IFN-γ pathway in myeloid cells restored migration features and collagen expression to control levels. Together, we identify a novel metastasis-promoting effect of Treg cells in the breast cancer microenvironment through regulation of ECM dynamics in an IFN-γ and myeloid cell-dependent manner in addition to their described effects on primary tumor growth.

## Results

### Treg cell ablation results in alteration of the ECM of primary mammary tumors

We have previously utilized the autochthonous MMTV-PyMT murine model of breast carcinogenesis crossed to the *Foxp3^DTR^* knock-in mouse model to investigate the effects of Treg cell ablation on breast cancer progression ^8,9,15^. To obtain an initial global characterization of the breast tumor matrix under control and Treg cell ablated conditions, we compared MMTV-PyMT tumors of at least 250mm^3^ in size obtained at several time points after targeting Treg cells with an acute diphtheria toxin (DT) treatment we previously shown to be transient, efficient and specific ^11,15,16^ (Fig. S1A-D). We harvested mammary tumors from control and Treg cell ablated mice at 4-, 10-, and 14-days after treatment, stained tumor sections using Masson trichrome (Fig. 1A-B) and quantified their collagen content (Fig. 1C). We observed a significant decrease in the amount of collagen per focal field in tumors devoid of Treg cells compared to control tumors at all time points analyzed (Fig. 1B-C). Of note, in the time frame analyzed (up to 14 days after DT injection), in this spontaneously developing model this treatment has no effect on tumor growth as measured by tumor volume ^10^ ^8^. Tumor associated collagen orientation is an important prognostic factor in breast cancer, with linearized fibers facilitating invasion and metastasis ^17,18^. We used multi-photon second harmonic generation (SHG) imaging and measured collagen fiber orientation in MMTV-PyMT tumor sections 4 days after Treg cell ablation. Interestingly, in addition to the reduction in collagen amounts, we also observed a small, but significant reduction of the linear, organized collagen structure in the samples devoid of Treg cells (Fig. 1D-E).

**Fig 1.**
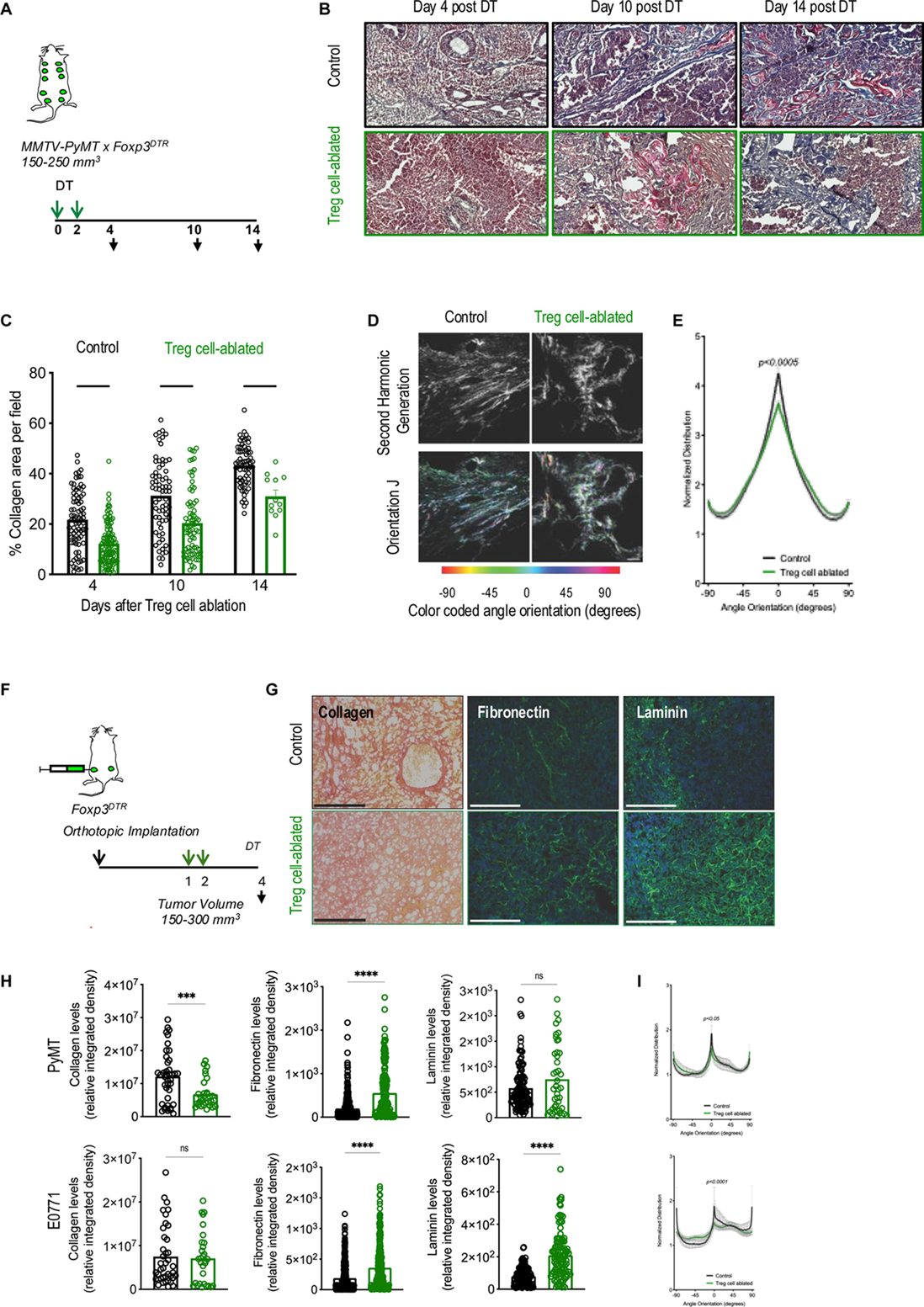
Treg cell ablation leads to dysregulation of extracellular matrix in breast cancer murine models. (A) Schematic of experimental design. Green arrows indicate diphtheria toxin (DT) treatment, black arrows indicate tissue collection. (B) Representative images of Masson’s trichrome staining of primary MMTV-PyMT tumors 4, 10, and 14 days post Treg cell ablation. Scale bar = 900 um. (C) Quantification of collagen-positive area in MMTV-PyMT tumors for control and Treg cell ablated animals, n = 2-4 mice per condition. 40-50 images at 10x magnification were analyzed per mouse using a combination of ImageJ and Masson’s trichrome de-convolution macro. Data is displayed as mean +/- SEM. **p<0.01 and ****p<0.0001. Statistical analysis was performed by two-way analysis of variance with Bonferroni multiple comparisons post-test. (D) Representative Second Harmonic Generation (SHG) images of Control and Treg cell ablated tumors. Scale bar = 100 um. (E) OrientationJ distribution output obtained from SHG images of Control and Treg cell ablated MMTV-PyMT tumors, n[=[5 tumors per group. Data is presented as mean values +/- SEM. Statistical analysis was performed by unpaired two-tailed Kolmogorov-Smirnov test with a 95% confidence level. (F) Schematic of experimental design. Black downward arrow indicates orthotopic implantation and tissue collection, green arrows indicate DT treatment, carried out at tumor volumes ∼150-300mm3 (G) Representative images of Picrosirius Red staining, fibronectin and laminin immunofluorescence staining (with DAPI) of primary orthotopic PyMT and E0771 tumors 4 days post Treg cell ablation or control conditions, in black and green color, respectively. Scale bar = 250 um. (H) Quantification of collagen-positive area (left), fibronectin (middle) and laminin (right) in PyMT and E0771 orthotopic tumors, respectively. n = 2-3 mice per cell line condition, bilateral orthotopic tumors. Analysis was performed using ImageJ. Data is displayed as mean +/- SEM. ***p<0.001 and ****p<0.0001. Statistical analysis was performed by unpaired t-test with Welch’s correction. (I) OrientationJ distribution output obtained from SHG images of Control and Treg cell ablated PyMT (top) and E0771 (bottom) tumors, n[=[2-3 tumors per group. Data is presented as mean values +/- SEM. Statistical analysis was performed by unpaired two-tailed Kolmogorov-Smirnov test with a 95% confidence level.

To explore the tumor ECM in more detail, we took advantage of the more synchronous tumor growth behavior and shorter timeline of our thoroughly characterized breast cancer orthotopic models, PyMT and EO771 cell lines. We utilized Picrosirius red staining to reveal collagen fibers in orthotopic primary mammary tumors from control and Treg cell ablated mice. In addition, we used immunofluorescence to detect expression of fibronectin and laminins, other major mammary gland ECM proteins associated with breast cancer and disease progression ^19^ (Fig. 1F). We collected tumors 4 days after DT treatment, time point at which we previously characterized triggering events without any effect on primary tumor growth, which begin to manifest 10 days after treatment ^11^. Consistent with the changes observed in the spontaneous MMTV-PyMT tumors, we detected reduced staining of collagen among Treg cell ablated PyMT-injected tumors compared to control (Fig. 1G-H, left panel), although this difference was not significant in E0771 orthotopic tumors (Fig. 1G-H, left panel). Yet, linear organization of collagen fibers was reduced in both PyMT and E0771 orthotopic tumors upon Treg cell ablation (Fig 1I, top and bottom, respectively).

Furthermore, fibronectin amounts were increased under Treg cell ablation dependent conditions in both cell models (Fig. 1G-H, middle panels). Laminin amounts were unchanged in PyMT tumors (Fig. 1G-H, right panel) but significantly increased in EO771 tumors upon Treg cell ablation (Fig. 1G-H, right panel). To evaluate the possibility of ECM changes being caused by systemic inflammation in a non-specific manner, we induced sterile systemic inflammation by intravenous delivery of LPS (800ug/kg for 4 days) and performed Picrosirius red staining of tumors (Fig. S2A). As expected, significant inflammation was observed (Fig. S2B-C), including previously reported increase in Treg cells ^20^. However, in this case collagen amounts in tumor ECM was increased rather than reduced (Fig. S2D), supporting a positive correlation with Treg cell presence. Overall, our data demonstrate significant ECM changes upon targeting the Treg cell compartment in advanced mammary gland tumors, although with tumor model-specific nuances.

### Decellularized ECM from Treg cell ablated tumors alters cell spreading and expression of EMT markers in tumor cells

The observed ECM alterations in murine mammary tumors devoid of Treg cells were significant, particularly of collagens, although complex and with model-dependent nuances. Instead of focusing on single changes, we reasoned that the remodeled matrix effects on the tumor cells would be the consequence of the combined global changes, therefore we decided to utilize the whole decellularized extract for subsequent functional experiments. Given the prominent role of the ECM on cellular processes leading to migration and invasion, we designed a battery of assays to evaluate this function.

To obtain cell-free ECM, we decellularized tumors following established protocols ^12,21^, and used this decellularized ECM (dECM) to embed tumor clusters or to coat tissue culture plates, as depicted in Fig. 2A. First, we resorted to a modified 3D microfluidic system to evaluate collective movement of cells seeded on different substrates in response to a gradient of SDF-1 in real time using confocal microscopy ^22^. In the original set up, tumor clusters are encapsulated in collagen I and loaded in the middle chamber of a microfluidic device ^22^. Here, PyMT- or EO771-derived tumor clusters were encapsulated in 3D custom dECM gels from matching tumors before loading in the device and allowed to settle on the gel for 48 hours while regular medium was delivered through the top and bottom fluidic lines (Fig. 2A). A static image was obtained at that time point (Fig. 2B), and tumor organoid area and perimeter were obtained as a measure of initial tumor spreading (Fig. 2C). Interestingly, both parameters were reduced in PyMT and EO771 organoids seeded on their respective dECM product from the Treg cell ablated conditions, indicating substrate-depending differences in the ability of the cells to spread (Fig. 2C, left and right panels, respectively). Furthermore, the collagen orientation index in these 3D chips was significantly reduced (Fig. 2D-E), as previously observed in spontaneous MMTV-PyMT (Fig. 1E) and orthotopic tissue sections (Fig. 1I). Importantly, we did not observe changes in the frequency of phospho-Histone 3^+^ proliferating cells in the tumor clusters in a period of 48hs, suggesting that the differences in their perimeter and area are not due to cancer cell growth (Fig. S3A-C). As reduced spreading might be the result of impaired epithelial-mesenchymal transition (EMT), the process that kicks off cancer cell dissemination ^23^, we ran a parallel experiment in which individual tumor cells were plated at sub-confluent levels on tissue culture plates pre-coated with dECM substrates using a thin layer method (Fig. 2A, bottom). This coating method was effective and reproducible, with no remaining cells observed in the dECM product (Fig. S4A-B) and minimal variation in total protein attached to each plate (Fig. S4D-E). After 24 hours of seeding, we harvested RNA and evaluated ECM-induced changes in four transcription factors known to induce EMT in breast cancer, namely Twist, Snail, Slug and Zeb ^24^. Western blot analysis of our cell lines showed that all these transcription factors are expressed at the protein level in both PyMT and EO771 to some degree, with exception of Slug, which was almost absent from PyMT cells (Fig. S5). Tumor cells that were co-cultured with dECM extracts from Treg cell ablated tumors showed reduced expression of the EMT transcription factor Snail, compared to those co-cultured with dECM extracts from control tumors (Fig. 2F). This was evident in both PyMT and EO771 murine cell lines. Additionally, Zeb and Twist levels were also reduced in PyMT cells when co-cultured with Treg cell ablated tumor dECM extracts compared to their control tumor dECM counterparts (Fig. 2F). Combined, these results suggest that Treg cell ablation-induced changes in the ECM lead to reduced ability of the tumor cells to undergo EMT.

**Fig 2.**
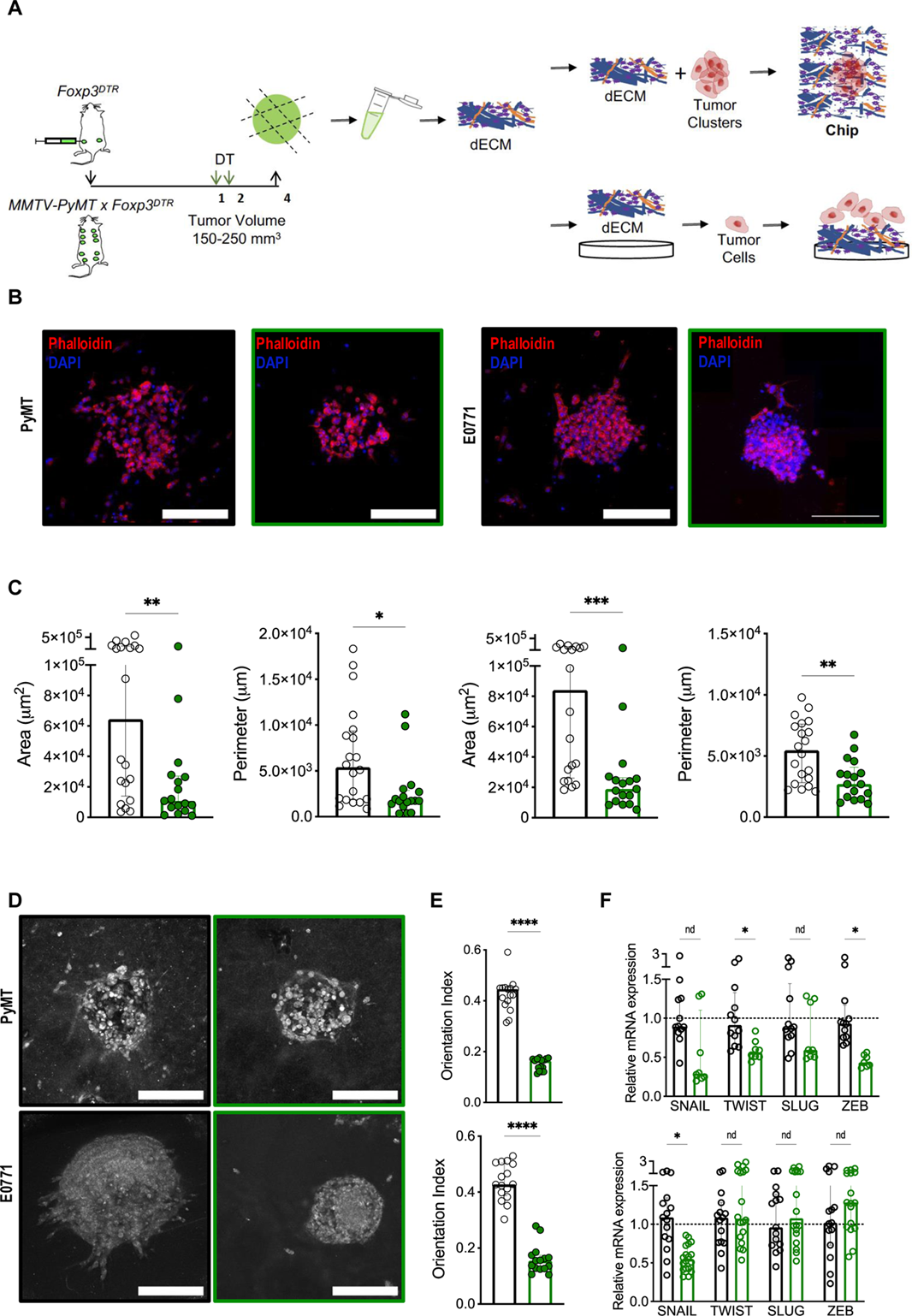
Treg cell ablation alters matrix fiber orientation and tumor cluster cell spread potential. (A) Schematic of experimental design. Black downward arrow indicates orthotopic implantation, green arrows indicate DT treatment, black upward arrow indicates tissue collection. Orthotopic tumors from control or Treg cell ablated conditions were processed to generate decellularized ECM (dECM) matrices. dECMs were used to mix with tumor organoids and load into the microfluidic device, or to coat multi-well plates for the for the plating of tumor cells for RNA isolation (B) Representative immunofluorescence images of PyMT (left) and E0771 (right) tumor clusters encapsulated in PyMT (or E0771) control or Treg cell ablated dECM, respectively. Phalloidin=red, DAPI=blue; scale bar = 250 um. (C) Quantification of tumor cluster area and perimeter of PyMT (left) and E0771 (right) after 2 days in culture. *n=3-4* technical replicates. Statistical analysis was performed by Mann-Whitney tests. (D) Representative collagen orientation images of PyMT (top) and E0771 (bottom) tumor clusters encapsulated in PyMT (or E0771 respectively) control or Treg cell ablated dECM. (E) Quantification of matrix fiber orientation (orientation index) in the static chip. Collagen=white; scale bar = 250 um. Data is shown as median +/- interquartile range. Statistical analysis was performed by Mann-Whitney tests. (F) Relative expression of EMT transcription factors Snail, Twist, Slug, and Zeb after 24 hours of tumor cell culture on their respective dECM coated plates. Triplicate analysis of PyMT (top) or E0771 (bottom), *n=2-3* animals per group. Data is shown as median +/- interquartile range. *p<0.05, **p<0.01, ***p<0.001, ****p<0.0001. Statistical analysis was performed by multiple Mann-Whitney tests.

### Treg cell-dependent changes in ECM alter tumor cell collective migration

To evaluate collective movement, after 48 hours of stabilization into the chip device, a SDF1 gradient shown to promote directional collective migration of tumors cell clusters in this system ^22^ was initiated through the microfluidics channel to induce tumor cell migration (Fig. 3A). As expected, PyMT- and EO771-derived tumor clusters encapsulated in control dECM responded to and migrated in the direction of the chemokine gradient (Fig. 3B-D and Videos S1 and S2). Importantly, similar tumor clusters embedded in Treg cell ablated dECM were profoundly impaired in their ability to migrate collectively following the gradient (Fig. 3B-D and Videos S3 and S4). Both migration efficiency and average velocity of PyMT or EO771-derived tumor cell clusters were severely reduced under similar conditions only differing on the source of the dECM (Fig. 3D). Together, our static dECM 3D co-culture and hydrogel, and the dynamic 3D microfluidic system results evidenced a significant loss of migration ability when cancer cells are encapsulated on Treg cell ablated tumor ECM. Our experiments suggest that Treg cells play a role in determining the chemical and structural composition of tumor-associated ECM, which in turn contributes to cell migration.

**Fig 3.**
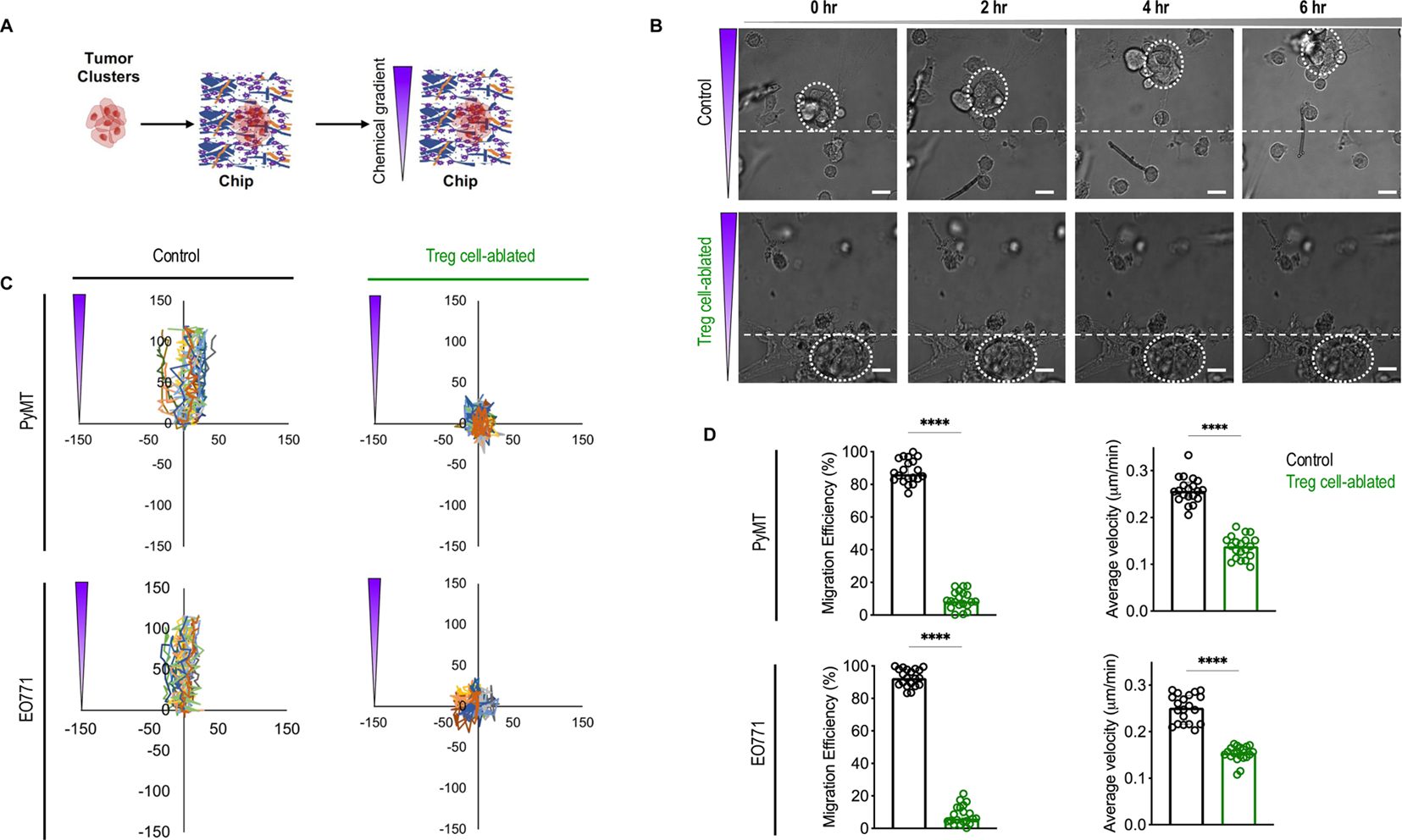
Treg cell ablation alters collective migration potential of primary tumor cell clusters. (A) Experimental schematic. dECMs hydrogels and tumor organoids were loaded into 3D microfluidics devices, cultured for 48 hours before inducing a biochemical gradient. (B) Representative brightfield images of live-cell imaging of PyMT tumor clusters embedded control (top) or Treg cell ablated (bottom) dECM in response to SDF1 chemokine gradient running from top to bottom (scale bar = 50 mm). (C) Migration maps of PyMT (top) or EO771 (bottom) tumor cell clusters in control or Treg cell ablated dECM. (D) Migration efficiency and average velocity quantification. Analyzed using Nikon Imaging Software and MATLAB. Data are represented as median +/- interquartile range. ****p<0.0001. Statistical analysis was performed using unpaired t-tests. *n=15-20* organoids from at least 3 different mice (∼4-8 organoids per mouse).

### Treg cell ablation leads to reduced circulating tumor cells and lung metastatic burden

So far, our *ex vivo* analyses demonstrate that Treg cell-dependent modifications of the ECM induce changes in EMT transcription factor expression and migration in response to known inducers of collective migration. These observations suggest that Treg cell-dependent matrix modifications may facilitate metastatic dissemination. To explore this biological process *in vivo*, we enumerated CD45^-^ pan-cytokeratin^+^ circulating tumor cells (CTCs) in the blood of control and Treg cell ablated mice using flow cytometry (Fig. S6), as indicated in the schematic (Fig. 4A). First, we assayed spontaneous MMTV-PyMT mice at several points after Treg cell ablation and found that the ratio of CTCs between control and Treg cell ablated tumors became increasingly more pronounced starting 4 days after Treg cell ablation, and statistically different 14 days after treatment (Fig. 4B). We then enumerated CTCs in PyMT and EO771 orthotopic models 4 days after Treg cell ablation (Fig. 4C). In this transplantable setting, the ratio of CTCs in the blood of control/Treg cell ablated mice was also significantly increased (Fig. 4D). Of note, at the evaluated time points (up to 14 days after treatment in spontaneous tumors and 4 days after treatment in orthotopic tumors), Treg cell ablation does not yet have any discernible effect on primary tumor size ^10,25^, indicating that primary tumor size is not responsible for number of tumor cells shed into the circulation ^26^.

**Fig 4.**
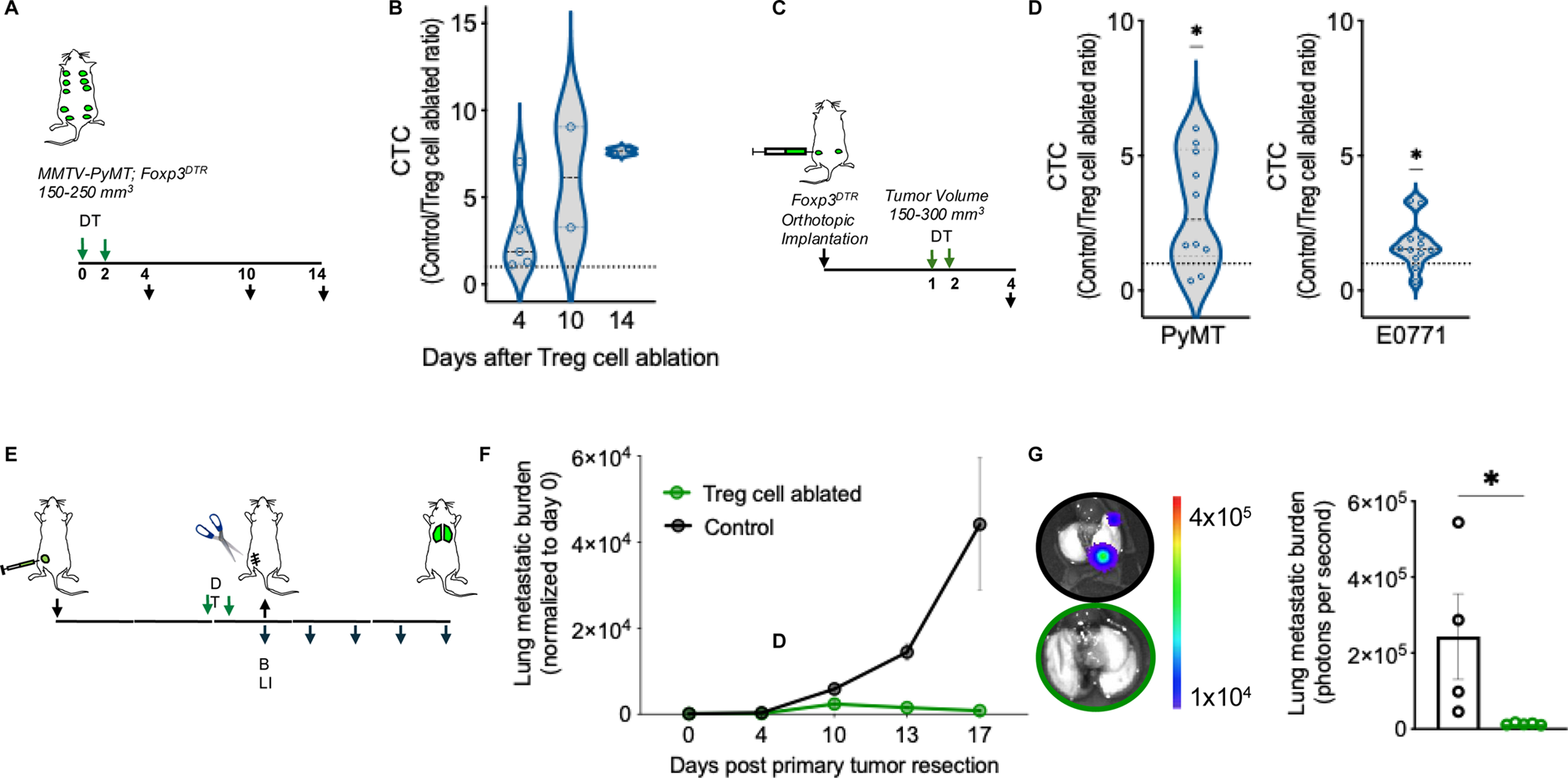
Treg cell ablation results in reduced amounts of circulating tumor cells in peripheral blood and reduced lung metastatic outgrowth upon neoadjuvant. (A) Schematic of experimental design for circulating tumor cell (CTC) enumeration, using spontaneous (A) and orthotopic (C) murine models. Black downward arrow indicates orthotopic implantation, green arrows indicate DT treatment, red arrow indicates CTC collection via cardiac puncture. (B) CTCs quantification of orthotopic tumor bearing mice as ratio between control and DT treated size-matched tumors, in a time-course manner and normalized to the amount of blood collected using beads. *n=2-4* mice per condition, per time-point. Data is shown as individual match-paired points, representative of one experimental set-up. Data is shown as median +/- SEM. ***p<0.001. Statistical analysis was performed by one-sample Wilcoxon signed rank test, with theoretical median set at 1.0. (D) CTCs quantification of orthotopic PyMT (left) and E0771 (right) tumor bearing mice as ratio between control and DT treated size-matched tumors, *n=10*-12 mice per condition. *p<0.05. Statistical analysis was performed by one-sample Wilcoxon signed rank test, with theoretical median set at 1.0. Data is shown as individual match-paired points. (E) Schematic of experimental design for lung metastatic neoadjuvant treatment. Black downward arrow indicates orthotopic implantation, green arrows indicate DT treatment, black upward arrow indicates tumor resection. IVIS imaging was performed post-tumor resection to monitor lung metastatic burden. n= 4-5 mice per condition. (F) Lung metastatic burden as photon flux shown as median +/- interquartile range. *p<0.05. Statistical analysis was performed by multiple Mann-Whitney tests. (G) Ex vivo lung tumor burden at the endpoint from experiment shown in (F), representative images (left) and photon flux quantification (right). Statistical analysis was performed by Mann-Whitney test. *p<0.05.

Reduced number of disseminated cells should ultimately diminish metastatic burden, as fewer cells would be able to seed secondary organs. To investigate this question, we employed a tumor resection protocol followed by longitudinal assessment of lung metastatic growth (Fig. 4E) that we have previously developed ^27^. Tumor-bearing mice were treated with DT to ablate Treg cells when tumors reached 250 mm^3^, and primary tumors were resected 4 days after treatment to prevent further dissemination, as per our previous observations (Fig. 4D). Lung metastatic burden was quantified by bioluminescence imaging (BLI) over the course of 2 weeks after resection, when all mice were sacrificed and lungs were imaged *ex vivo* (Fig. 4E). As expected, control mice started developing BLI-detectable lung metastasis from about a week after tumor resection, and continue growing rapidly (Fig. 4F, black line). On the contrary, animals that had been subjected to Treg cell ablation before tumor resection had significant reduction in lung metastatic growth (Fig. 4F, clover line) and lung metastatic burden at the end of the experiment, compared to their control counterparts (Fig. 4G). Overall, these findings suggest that the defects in migratory properties observed *ex vivo* when tumor cells were seeded on dECM from Treg cell-ablated conditions also happen *in vivo*, where less tumor cells can disseminate and contribute to establish lung metastasis.

### Treg cell-driven ECM structural and functional changes requires IFN-***_γ_***

Our previous work on Treg cell driven primary tumor growth uncovered a critical role for Treg cell suppression of IFN-γ production by CD4T cells in the breast tumor promoting function of inflammatory monocytes and tumor associated macrophages. Therefore, to explore whether IFN-γ also plays a role in the ECM changes we identified, we utilized loss of function approaches. First, we quantified collagen fibers by Picrosirius red staining in orthotopic PyMT tumors of mice in which we ablated Treg cells in the presence of neutralizing antibodies against IFN-γ or with genetic deletion of this cytokine (Fig.5 A). We observed that the reduction in collagen staining resulting from Treg cell ablation is significantly reversed if IFN-γ is neutralized or absent (Fig.5B-C), whereas interfering with IFN-γ alone has no significant effect on the collagen matrix. Interesting, lysates from these tumors have an increased collagenase activity only in Treg cell ablated conditions. While this change is not statistically significant, it shows a noteworthy inverse correlation with the Picrosirius red assessment (Fig. S7).

**Fig 5.**
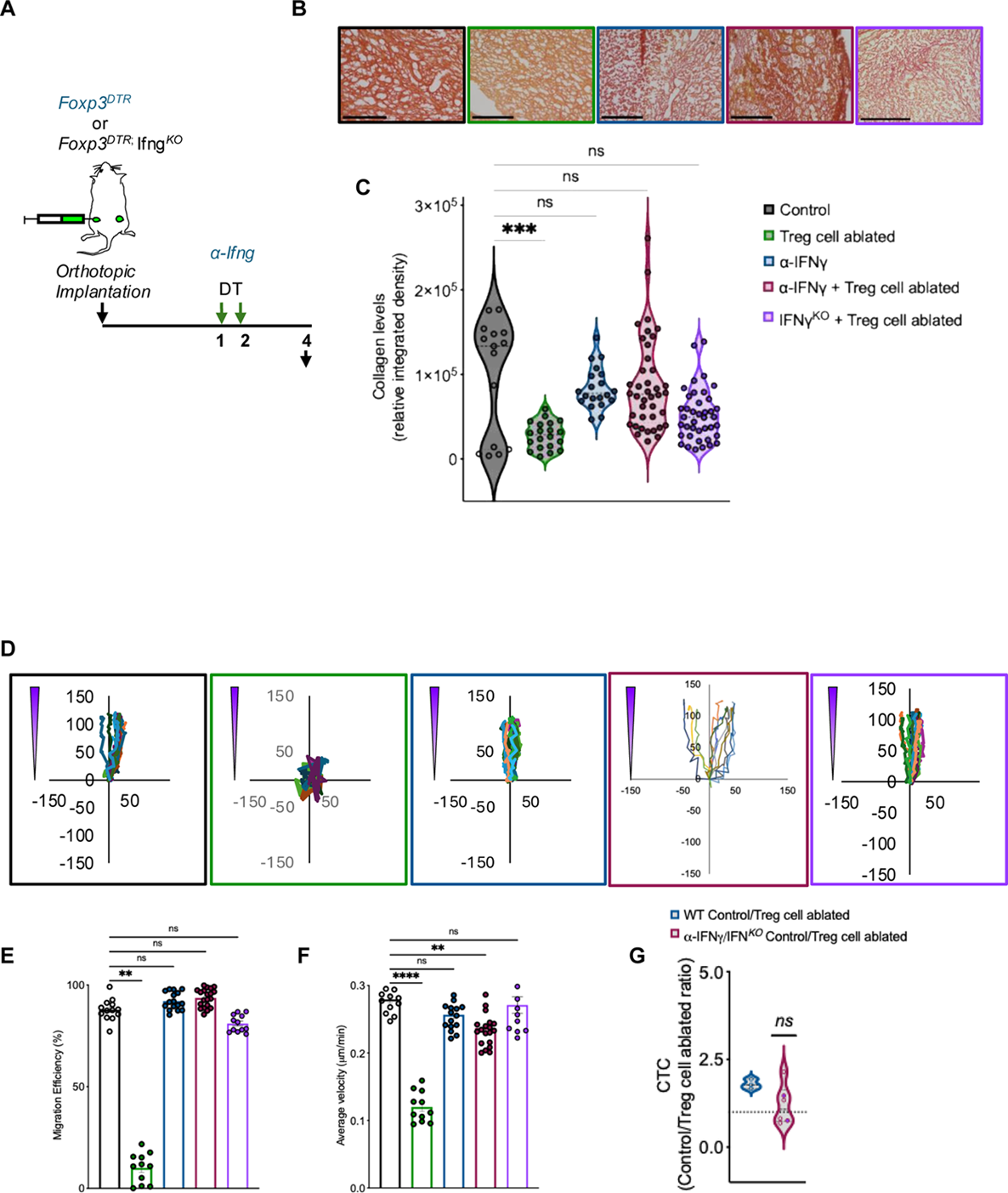
Treg cell ablation-dependent ECM remodeling phenotypes are IFN-_γ_-dependent. (A) Schematic of experimental design for blocking IFN-γ, by either neutralizing antibody (a- IFN-γ) or genetic model (*IFN-*_γ_*KO* Foxp3^DTR-GFP^). Black downward arrow indicates orthotopic implantation, green arrows indicate DT treatment, red downward arrow indicates anti- IFN-γ treatment (if applicable), black upward arrow indicates tissue collection. (B) Representative images of Picrosirius Red staining of primary orthotopic PyMT tumors for control (black), Treg cell ablation (labeled as DT, green), neutralizing IFN-γ antibodies (labeled as a- IFN-γ, red), a- IFN-γ +DT (blue), and *IFN-*_γ_*KO*+DT (orange) conditions. Scale bar = 250 um. (C) Quantification of collagen-positive area in PyMT orthotopic tumors. n = 2-4 mice per condition. Analyzed one bilateral orthotopic tumor, per mouse. 10 representative ROIs were selected from Keyence-generated stitched images at 20x magnification, per bilateral orthotopic tumor. Analyzed using ImageJ. Data is displayed as mean +/- SEM. ***p<0.001. Statistical analysis was performed by Kruskal-Wallis test with Dunn’s multiple comparisons post-test. (D) Migration maps of PyMT tumor cell clusters in control, Treg cell ablated, a- IFN-γ, and a- IFN-γ +DT dECM. (E) Migration efficiency and average velocity quantification. Analyzed using Nikon Imaging Software and MATLAB. Data are represented as median +/- interquartile range. ****p<0.0001. Statistical analysis was performed using by Kruskal-Wallis test with Dunn’s multiple comparisons post-test. (F) CTC quantification of orthotopic tumor bearing mice as ratio between control and treated size-matched tumors, *n=2*-4 mice per condition. **p<0.01. Statistical analysis was performed by one-sample Wilcoxon signed rank test, with theoretical median set at 1.0. Data is shown as individual match-paired points.

Next, we obtained decellularized ECM from the same experimental set up to assess the consequences of IFN-γ loss of function in the tumor organoid migration phenotype in the 3D chip assays. As expected, tumor clusters encapsulated in Treg cell-ablated dECM showed reduced migration ability and speed compared to those encapsulated in control dECM (Fig 5D-E). However, tumor clusters encapsulated in an IFN-γ -deprived dECM, or those in the dECM originated from tumors in which IFN-γ and Treg cells were simultaneously ablated, showed similar migratory ability as those encapsulated in control dECM (Fig 5D-F).

Lastly, we evaluated the effect of IFN-γ blockade in tumor cell dissemination by enumerating CTCs. As in previous experiments ratio of CTCs between control and Treg cell-ablated mice indicated reduced dissemination in the absence of Treg cells. However, similar number of CTCs were found in control and Treg cell ablated conditions with simultaneous blockade of IFN-γ (1:1 ratio), suggesting that the reduction in tumor cell dissemination from tumors mediated by Treg cell targeting requires IFN-γ (Fig. 5G).

Altogether these experiments indicate that structural and functional changes in the ECM observed upon Treg cell targeting are mediated through IFN-γ.

### Treg cell-dependent matrisome signature correlates with long-term survival in human breast cancer

An ensemble of genes that encode ECM and ECM-modifying proteins called the matrisome has been assembled bioinformatically based on proteomics analysis of various tumor ECMs ^28-30^. Since our findings demonstrated a functional effect of Treg cell-dependent remodeling of the ECM and its influence over tumor cell dissemination, we aimed at exploring the Treg cell-dependent regulation of the matrisome in our models using bulk RNA-sequencing from control and Treg cell-ablated EO771 tumors. Initial unsupervised clustering analysis revealed distinct treatment-dependent clusters in the PCA plot, except for one outlier which was subsequently excluded from the analysis (Fig. S8A). We compared control and Treg cell ablated transcriptomic changes and queried the status of the 1110 matrisome genes to identify significantly distinct changes in core, affiliated and regulatory ECM proteins. This analysis revealed a group of 73 ECM-associated differentially expressed genes (DEGs), defined by an FDR <= 0.05, and a 1.4 absolute log2 fold change among Treg cell ablated tumors compared to control tumors (Fig. 6A and Table 1). Of those, 40 were up-regulated and 33 were down-regulated.

**Fig 6.**
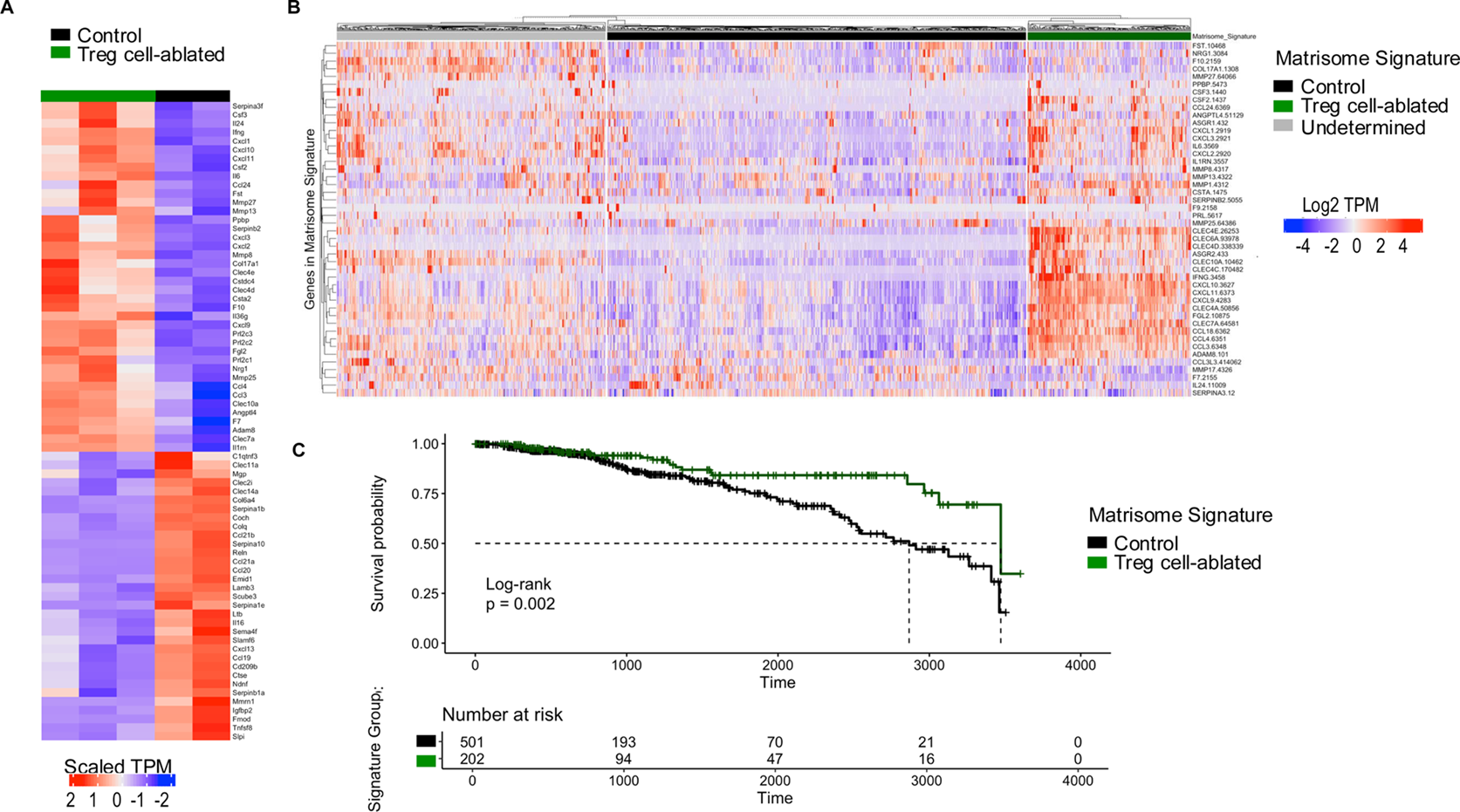
Treg cell ablated matrisome gene expression from EO771 model correlates with long-term survival in TCGA breast cancer cohort. (A) Heatmap of matrisome genes that were significantly differentially expressed (DEGs, >1.4 log2 fold change, FDR<0.05) between Treg cell ablated (green lines) and control (black lines) murine tumors. (B) Heatmap of K-means clustering (k=3) with Euclidean distance among 1079 TCGA breast cancer patients using human orthologs for the 40 upregulated matrisome DEGs. Clusters are labeled as Control-like (low expression), Treg cell ablated-like (high expression), or Undetermined (top color bar) based on the similarity of the human gene expression to that of the mouse matrisome gene signature. (C) Kaplan-Meier survival analysis among clustered “Treg cell ablated-like” and “Control-like” TCGA breast cancer patients over a 10-year period using the 40-gene highly expressed matrisome signature.

**Table 1.**
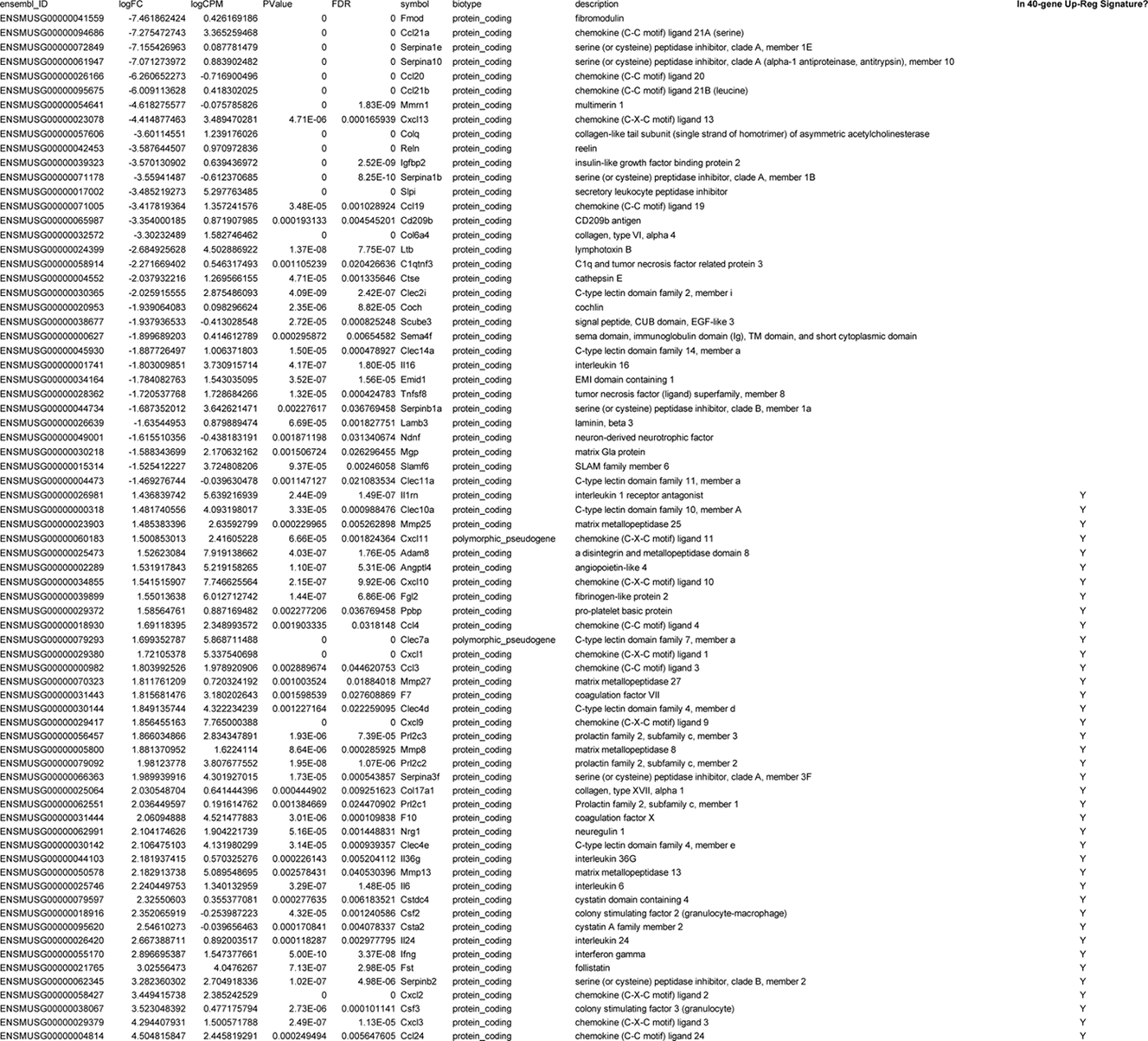
73 differentially expressed ECM-associated genes (DEGs. ). List of 73 DEGs defined by an FDR <= 0.05, and a 1.4 absolute log2 fold change among Treg cell ablated tumors compared to control tumors. Columns indicate ensemble ID, log fold change (logFC), log counts per million (logCPM), p-value (PValue), false discovery rate (FDR), gene symbol (symbol), and description.

In the past, we have shown that Treg cell-driven TAM transcriptomic signatures can be found in human breast cancer samples, and that tumors which harbor similar molecular signatures to those of Treg cell ablated TAMs have better survival ^11^. We wondered whether similar Treg cell-dependent matrisome signatures were also conserved in human breast cancer samples, and whether they had any correlation with patient outcome. We utilized the 46 human orthologs corresponding to the 40 upregulated matrisome DEGs to cluster breast cancer patients in the TCGA cohort into “Control-like” and “Treg cell ablated-like” groups, using k-means clustering with Euclidean distance (Fig. 6B). Additionally, we performed Kaplan-Meier survival analysis among these two groups, which revealed that the Treg cell-ablated-like group displayed an increase overall survival compared to patients with the “Control-like” matrisome signature (Fig. 6C). Interestingly, this increase in overall survival was observed in a 10-year period analysis but not yet manifested in a standard 5-year period analysis (Fig. S8B-C). As breast cancer diagnosis is often followed by a period of latency before symptomatic metastatic disease manifests ^31^, this delayed survival benefit observed in patients with Treg cell ablation-like matrisome signature is consistent with the reduced tumor cell dissemination phenotype we observe in mice, consistent with seeding a reduced number of DTCs that may result in reduced lung metastatic outgrowth at a future time. Together, these findings suggest that our observations in breast cancer murine models may be relevant to breast cancer disease in humans.

### Matrisome DEG signature is highly upregulated among tumor macrophages

Cancer-associated fibroblast deposition and remodeling of the ECM is widely recognized as critical contributor to breast cancer progression ^32^. However, many other cell types in the TME can contribute to the remodeling, including tumor cells, myeloid cells, endothelial cells. Recently, TAMs have been proposed to both deposit ^12,14^ or remodel ^33^ the ECM. To investigate this aspect globally, we performed single-cell RNA seq (scRNA-seq) analysis on control and Treg cell-ablated breast primary tumors. K-means clustering was used to establish different clusters of cell types, and UMAP (Uniform manifold approximation and projection) was used to visualize these clusters (Fig 7A). Cell type clusters were defined by identity marker genes (Fig. S9) and by GSEA of top DEGs from each cell cluster. A broad range of TME cells were represented: B cells (14%), T cells (30%), NK/NKT cells (3%), macrophages (8%), tumor cells (13%), fibroblast cells (25%), erythroid cells (6%), and granulocytes (1%) (Fig 7A). To identify which cell type (or cell types) might be responsible for the functional reprogramming of the ECM, we calculated an ECM signature score based on the previously described matrisome DEG signature (Fig. 6A and Table 1). This ECM signature score was overlaid onto the UMAP of cell clusters (Fig 7B). Interestingly, we observed the most elevated ECM signature scores in the macrophage cluster, compared to the other TME cell types (Fig 7B). In addition, fibroblast and tumor cell clusters also displayed somewhat elevated, but heterogeneous expression of the ECM signature scores (Fig 7B).

**Fig 7.**
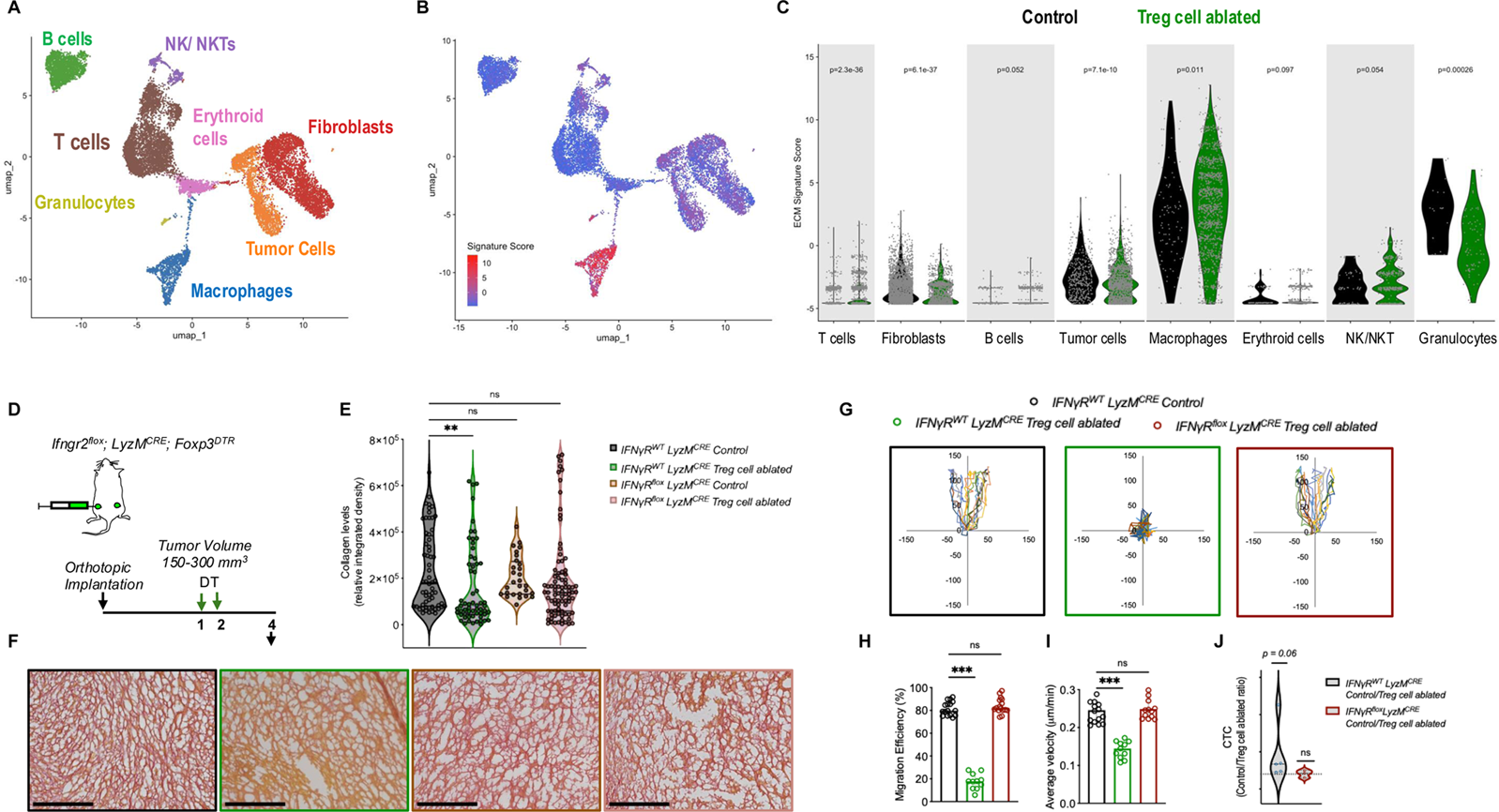
Treg cell ablation-dependent ECM remodeling phenotypes require IFN-_γ_ signaling in TAMs. (A) K-means clustering of good quality, viable cells from 4 primary mammary tumors, visualized as a UMAP (Uniform manifold approximation and projection) plot (left). Different cell type identities are color coded and labelled accordingly. UMAP colored by level of ECM signature scores, which reflects level of expression for ECM genes identified using the bulk RNASeq data (right). (B) Violin plots for level of ECM signature scores, by each cell cluster type and treatment group (control in black, Treg cell-ablated in green). (C) Schematic of experimental design for blocking IFN-γ in macrophages using a genetic model (*IFN-*_γ_*R2^flox^ LysM^CRE^ Foxp3^DTR-GFP^*). Black downward arrow indicates orthotopic implantation, green arrows indicate DT treatment, black upward arrow indicates tissue collection. Representative images of Picrosirius Red staining of primary orthotopic PyMT tumors for wildtype-control (black), wildtype-Treg cell ablated (green), flox-control (purple), flox-Treg cell ablated (pink) conditions. Scale bar = 250 um. (D) Quantification of collagen-positive area in PyMT orthotopic tumors. n = 4-5 mice per condition. Analyzed one bilateral orthotopic tumor, per mouse. 10 representative ROIs were selected from Keyence-generated stitched images at 20x magnification, per bilateral orthotopic tumor. Analyzed using ImageJ. Data is displayed as mean +/- SEM. **p<0.01. Statistical analysis was performed by Kruskal-Wallis test with Dunn’s multiple comparisons post-test. (E) Migration maps of PyMT tumor cell clusters in wildtype-control (black), wildtype-Treg cell ablated (green), and flox-Treg cell ablated dECM (pink). (F) Migration efficiency and average velocity quantification. Analyzed using Nikon Imaging Software and MATLAB. Data are represented as median +/- interquartile range. ***p<0.001. Statistical analysis was performed using by Kruskal-Wallis test with Dunn’s multiple comparisons post-test. (G) CTC quantification of orthotopic tumor bearing mice as ratio between control and Treg-cell ablated size-matched tumors for wildtype and flox mice, *n=3-5* mice per condition. Statistical analysis was performed by one-sample Wilcoxon signed rank test, with theoretical median set at 1.0. Data is shown as individual match-paired points.

Next, we compared ECM signature scores between control and Treg cell-ablated conditions for each cell type (Fig 7C). Importantly, we observed that the ECM signature scores were only significantly elevated upon Treg cell ablation in the macrophage compartment but not fibroblasts or tumor cells, where a small but significant downregulation of the score was observed (Fig 7C). A similar trend was observed in the granulocyte cluster; however, our processing pipeline only allowed for a miniscule representation of the granulocyte compartment, with less than 1% of total cells, making this observation inconclusive. Overall, these results point to an important role of macrophages in the ECM remodeling process.

To specifically investigate whether our observed ECM changes could be explained by IFN-γ - mediated TAMs reprogramming, we utilized IFN-γR2*^flox^*LyzM*^CRE^* Foxp3*^DTR-GFP^* mice (Fig 7D), in which macrophage-specific IFN-γ -signaling is absent.

First, we used this strain to implant orthotopic PyMT tumors, ablate Treg cells and assess the amount of collagen fibers by Picrosirius red staining (Fig. 7E-F). As expected, Treg cell ablation resulted in significant reduction of collagen staining compared to control in wild type mice (Fig 7E-F), however in mice with deficiency of IFNγR in the myeloid cell compartment, Treg cell ablation collagen fibers were similar than control mice (Fig 7E-F).

Next, we assessed the effect of dECM obtained from these mice in the tumor fragment migratory behavior. As in our previous experiments, tumor clusters encapsulated in dECM from Treg cell-ablated mice showed reduced migratory ability and velocity, compared to those encapsulated in control dECM (Fig 7G-I). However, Treg cell ablation in mice where myeloid cells are insensitive to IFN-γ signaling showed similar migratory ability and velocity as those encapsulated in control wildtype dECM (Fig 7G-I).

Last, to determine if this macrophage IFN-γ -mediated effect holds true *in-vivo*, we assessed tumor cell dissemination by enumerating CTCs. As before, we calculated the ratio of blood CTCs in control/Treg cell-ablated wildtype and macrophage IFN-γ-insensitive mice (Fig 7J). Again, the CTC ratio of control/Treg cell-ablated was significantly higher than 1, indicating more tumor cell dissemination in control animals compared to Treg cell-ablated animals (Fig 7J). However, the ratio between CTCs in control/Treg cell-ablated conditions among macrophage IFN-γ -insensitive mice was close to 1, suggesting similar amounts of CTCs among these two experimental groups (Fig 7J), reverting towards control levels despite Treg cell-ablation. Altogether these experiments suggest that IFN-γ -signaling in macrophages is necessary to drive the Treg cell-dependent ECM-related phenotypes.

## Discussion

Overcoming the strong immune suppression of the breast tumor microenvironment is one of the main challenges for the development of successful immunotherapeutic modalities, as well as to improve regular chemotherapeutics.

It is well established that the tumor-associated ECM, the non-cellular tridimensional network in which the tumor resides, is a key regulator of many critical cancer processes, most notably metastatic dissemination. Moreover, an important emerging function of the ECM is that of immune modulation at virtually all levels of the anti-tumor immune response, especially the control of immune cell trafficking through mechanical forces, generation of matrikines or modulation of chemokines and the regulation of immune cell activation and polarization states ^34^. In recent years, newfound appreciation has developed for the impact that physiological or exogenous conditions might have on metastatic dissemination through their direct effects on the ECM. For example, obesity induces myofibroblast deposition of denser ECM and subsequent increased stiffness ^34^. In addition, obesity drives increased collagen IV in the breast cancer ECM ^35^, both these changes leading to increased metastatic dissemination. Intriguingly, a recent report demonstrates that traditional chemotherapy also induces remodeling of the ECM that results in increased tumor cell invasion, which might explain the disconnect between cytotoxicity and invasion in some breast cancer patients undergoing neoadjuvant chemotherapy ^36^.

The previously underappreciated crosstalk between the immune system and the matrix has regained attention as the highly dynamic nature of this interplay is revealed, particularly in the context of the ECM being a strong obstacle to overcome for immunotherapy ^37,38^. While the consequences of ECM remodeling on regulation of immunity are becoming clear, we pondered whether immune-targeted interventions may have unanticipated consequences on tumor evolution through their impact on the tumor-associated ECM.

Tumor Treg cells are a highly desirable target for immune intervention ^39,40^, but faces the formidable challenge of uncoupling its effects on the tumor from their role as enforcers of peripheral tolerance. Therefore, increasing our mechanistic understanding of their tumor-specific functions in the TME is critical to develop novel immunotherapeutic modalities. In this work, we use a combination of *in vivo* interventions with *ex vivo* cellular, molecular and novel bioengineering approaches to demonstrate that specific targeting of the Treg cells results in previously unrecognized changes to the ECM that ultimately impair the process of cell migration and metastatic dissemination. Efficient decellularization of the ECM from treated and untreated tumors demonstrates its direct effect on tumor cell behavior. This new observation complements the modulation of primary tumor growth by macrophages we previously described ^11^. Further, we now show that Treg cell-dependent changes in the matrisome ^28,29^ can identify a breast cancer patient population with delayed improved overall survival, where the survival advantage manifests only when assessed 10 years post-primary tumor excision. Interestingly, this is different than similar analysis we performed with a signature derived from Treg cell-specific changes in the TAM compartment, where survival benefit in patients that display the TAM signature was evident within a 5-year period ^11^. It is now widely demonstrated that dissemination of tumor cells in breast (and other) cancers occurs early, probably even before the primary tumor is diagnosed ^31^. But metastatic relapse frequently occurs with a significant latency of 10 to 15 years, often following a prolonged period of clinically undetected disease, or “dormancy” ^31,41^. The discrepancy we observe when using the matrisome versus the TAM-derived signatures might be a reflection of this biology: while TAMs can have multiple effects on the tumor, including rapid tumor cell killing, Treg cell-dependent modifications of the ECM result in impaired cell migration and reduced number of CTCs, an effect that will impact metastatic relapse later in the course of disease.

Our studies provide the framework for an important number of new questions to address. First, what are the specific changes in the ECM that drive tumor cell behavior. We have observed changes in collagen and collagen orientation, fibronectin and laminin in tumors devoid of Treg cells. Our studies demonstrate the global effect of the resulting ECM on cancer cells, however which specific molecules or structures drive this effect remains an important question to investigate. Mass spectrometry characterization of the ECM and fiber orientation studies in both tumor settings, followed by targeted approaches using traditional single substrate cell-ECM assays ^42-44^ should help clarify this important point. Furthermore, it will be valuable to determine whether those changes extend to other tumor types. The answer to these questions may inform mechanistic insights with therapeutic value and reveal predictive biomarkers for clinical prognosis^37,45^.

We demonstrate significant dependency of the ECM reprogramming on IFN-γ and IFN-γ- dependent myeloid signaling. Whether the TAMs directly remodel the ECM or modulate the activity of CAFs or other cells remains to be formally addressed. In our previous studies, we observed ECM and ECM remodeling genes as differentially expressed in TAMs upon Treg cell targeting (^11^, data not shown), and here we find TAMs displayed the most significant matrisome signature. Moreover, monocytes and TAMs have been previously shown to be important players in ECM remodeling ^12,14,33,46,47^. On the other hand, tumor cells and cancer associated fibroblasts (CAFs) are known to be the main responsible for tumor-associated ECM deposition and remodeling ^48,49^, and CAF matrix-remodeling transcriptional programs have been recently observed in lung cancer upon Treg cell ablation ^50^. Whether Treg cell effects on the ECM are mediated by one or several of these cellular types remains to be elucidated through genetic ablation or depletion of relevant cell populations and signaling pathways.

The last decade has provided a new framework for the understanding of Treg cell biology and established the existence of Treg cells resident in tissues (“tissue Treg cells”) that actively participate in tissue physiology and disease by engaging in functions conferred by sensing the specific microenvironment. An emergent theme is that tissue Treg cells are critical for repair mechanisms ^15,51,52^, so active engaging the tissue matrix would be fitting. As tumors are considered wounds that do not heal, this ability of tumor Treg cells would be advantageous in the TME.

Our work reveals a previously unrecognized Treg cell-dependent regulation of the ECM that contributes to metastatic dissemination. Importantly, by providing a tangible example of regulation of such a critical tissue component, it underscores a function of Treg cells that goes beyond their traditional immunological role and demonstrates their potential for the physiological regulation of tissue biology.

## Acknowledgements

We are grateful to C. Lemmon (VCU) and R. Heise (VCU) for invaluable advice. This work was supported by a Research Scholar Grant RSG-21-100-01-IBCD from the American Cancer Society to P.D.B. and a Theory Lab TLC-21-158-01 grant from the American Cancer Society to P.D.B. and P.Y.H.. P.D.B. was also supported by the Susan G. Komen Foundation CCR18548205, the V Foundation V2018-022, and the NCI R37 MERIT Award CA269249. P.Y.H. was supported by NSF CAREER Award 2145756. J.J.B.-C. was supported by NCI R01 CA244780, NCI R03 CA270679, NCI R61 CA278402, the Irma T. Hirschl Trust, the Emerging Leader Award from the Mark Foundation and Tisch Cancer Institute NIH Cancer Center grant P30 CA196521. A.L.O. was supported by CTSA award UL1TR002649 from the National Center for Advancing Translational Sciences. Services obtained through the VCU Massey Cancer Center Flow Cytometry, Cancer Mouse Models Shared Resource, Microscopy Shared Resource, and Tissue and Data Acquisition and Analysis Core (TDAAC) were supported, in part, with funding from NIH-NCI Cancer Center Support Grant P30 CA016059.

## AUTHOR CONTRIBUTION

A.D.G-S. and P.D.B. designed the project, analyzed the experiments, and wrote the manuscript. A.D.G-S, H.S., J.M.R., N.M.C., W.D., L.M.M., A.D.H., T.C. performed most of the experiments. J.Y.L and P.Y.H. performed the organ-on-a-chip experiments. J.B. and J.J.B.C. performed second harmonic generation analysis. M.G.D. performed the RNA sequencing pre-processing. A.L.O. performed the bioinformatics analysis of the human datasets and scRNASequencing analysis. P.D.B. supervised all aspects of the project. All authors read and commented on the manuscript.

## METHODS

### Study Design

Animal studies were conducted in concordance with Virginia Commonwealth University’s Division of Animal Research (DAR) and Institutional Animal Care and Use Committee (IACUC) approved protocol. Group sizes of each experiment were determined based on an 80% power of detecting a medium-to-large, standardized effect size 0.5-0.75 (≤5% chance of error) assuming 0.2 variability. Mouse subjects were randomized prior to treatments based on tumor volume as measured by caliper measurements, and each experiment was performed in triplicate, unless specified. Mice were monitored daily for health parameters, and they were euthanized at indicated tumor volumes within the limits permitted in our IACUC protocol.

### Experimental Models

C57BL/6 *MMTV-PyMT Foxp3^DTR-GFP^*, *Foxp3^DTR-GFP^ LysM^CRE^*, and *Foxp3^DTR-GFP^ IFN-*_γ_ *^KO^* mice were generated before ^15^. C57BL/6 *MMTV-PyMT Foxp3^DTR-GFP^*mice monitored bi-weekly for mammary primary tumors by caliper measurement of tumor length (L) and width (W). Tumor volumes were calculated by πLW^2^/6 until they reached at least 150 mm^3^, and mice were randomly grouped into two groups to initiate treatment. Luciferase-transduced PyMT ^8^ and EO771 cell lines (C3H Biosystems) were previously described, and cultured in RPMI-1640 (Cytiva Hyclone), supplemented with 10% fetal bovine serum (FBS, R&D Systems), 1% penicillin-streptomycin (Gemini BioProducts), and 0.5% amphotericin (Gemini BioProducts). Cell lines were orthotopically injected at a seeding density of 150,000 to 500,000 cells/injection at 1:1 with Matrigel Cultrex Basement Membrane Extract, Type 3 (R&D) into the fourth mammary gland of isoflurane-anesthetized *Foxp3^DTR-GFP^*mice in a bilateral manner. Mice were monitored twice a week until tumor became palpable, and caliper measurements were taken as before. Treg cell ablated group was treated with two doses of diphtheria toxin (DT, List Labs) at a 50ug/kg concentration, on two consecutive days (orthotopic tumors) or on days 0 and 2 (spontaneous tumors), as done before ^8,9^. Mice were humanely euthanized at end points indicated in the figures, blood collected by cardiac puncture and tissues were harvested. LPS treated group received four doses of Lipopolysaccharides (from Salmonella Minnesota, R595 Re TLRgrade™, VWR) at 800 μg/kg concentration, on four consecutive days. IFN-γ neutralization group received one dose of neutralizing antibodies against IFN-γ (clone XMG1.2) at 1mg, together with the second dose of DT, as done before^8^.

### Histological Analysis of Mammary Primary Tumors

Primary tumor tissues were collected at indicated endpoints. Tissues collected for histological analysis were fixed in 4% paraformaldehyde (PFA, Sigma Aldrich) for 24 hours at 4•C. For histochemistry stains, tissues were transferred to 70% ethanol (KOPTEC) and embedded in paraffin blocks. For immunofluorescence stains, tissues were transferred to 30% sucrose (Thermo Scientific) for 24 hours at 4•C, then 1:1 OCT (Fisher Healthcare) and 30% sucrose solution for 24 hours at 4•C on a rotator, then twice rinsed in OCT for 30 minutes on ice and finally embedded in OCT over dry ice to obtain blocks. 7uM thick tissue sections were cut and mounted on glass slides for staining.

### Histological Staining

NovaUltra IHC World Masson’s Trichrome staining kit (IHC World) and Picrosiurus Red (Direct Red 80, Sigma Aldrich; acetic acid, Fisher Chemicals; saturated aqueous solution of picric acid, Aristar) staining were used as per manufacturer’s instructions (https://www.ihcworld.com/_protocols/special_stains/masson_trichrome.htm) (https://www.ihcworld.com/_protocols/special_stains/sirius_red.htm). Slides were mounted with Permount mounting media (Electron Microscope). Stained Masson’s Trichrome sections were imaged with Zen2 software (blue edition, GmbH 2011) on a brightfield Zeiss Axio Imager microscope at 10X magnification. 40-50 images were taken per tumor, at the tumor edges. Analysis was performed on ImageJ, with the built-in Masson’s trichrome de-convolution macro.

Stained Picrosirius Red sections were imaged with BZ-X800 Viewer software on a Keyence BZ-X800 microscope at a 20X magnification. The entire tumor was imaged and stitched together using the Keyence proprietary software. 10 representative ROIs were selected from each image and analyzed on ImageJ, using an adapted macro from ImageJ’s “Quantifying Stained Liver Tissue”, (https://imagej.nih.gov/ij/docs/examples/stained-sections/index.html).

### Immunofluorescence staining

Frozen sections were thawed at room temperature (RT) overnight and rinsed in 1X PBS (Quality Biologicals). Slides were blocked with 5% BSA (Fisher Bioreagents), 1% normal goat serum (Sigma Aldrich) in 1X PBS for 1 hour at room temperature. Slides were washed in 1X PBS, and incubated with primary antibodies resuspended in 0.5% BSA, overnight at 4C. Slides were washed in 1X PBS and incubated with secondary antibodies for 1 hour at RT in a humidity chamber, covered. Slides were washed with 1X PBS and then mounted with Vectashield Hard Set with DAPI (Invitrogen). Fibronectin was stained using a rabbit polyclonal antibody (Abcam Cat#ab2413, 1:500), followed by anti-rabbit Alexa Fluor 488 conjugated secondary antibodies (Invitrogen Cat#A11008, 1:1000). Laminin was stained with a rabbit polyclonal antibody (Abcam Cat#ab11575; 1:500), and same secondary antibody.

Stained sections were imaged on a Keyence BZ-X800, using GFP filter cube (Keyence) at 20X magnification. The entire tumor was imaged and stitched together using the Keyence proprietary software. Images from the tumor edges were analyzed using ImageJ with custom macros and normalized to tissue area. Cells in static hydrogels were fixed in 4% PFA and blocked in 1X PBS with 1% BSA and 0.1% Tween20 (Bio-Rad). Cells were washed in PBS with 1% Tween20. After fixing and blocking, cells were stained for phalloidin 568 (Invitrogen, Cat#A12380, 1:500) and nuclei (DAPI, Thermo Fisher, Cat#D1306). Imaging was performed with ZEN 3.3 software using a Zeiss LSM980 confocal microscopy (Zeiss).

### Second harmonic generation image acquisition and analysis

Images were acquired with Olympus FluoView software (FV-10-ASW 3.1) on an Olympus FV1000MPE multiphoton microscope (Olympus) equipped with a Coherent Chameleon Vision II laser, tunable from 680 nm to 1,080 nm; an excitation line of 880 nm was used for Second Harmonic Generation (SHG) using a 25x/1.05NA. Frame size was set to 512X512 pixels (X/Y) for a final lateral pixel size of 0.9944 mm, and images were acquired at 8 bits. The acquisition was performed at line sequential scan mode, with a pixel dwell time of 12.5 ms/pixel, and a frame average of 2. For z-stacks, the z-step size was set at half the axial resolution (5 mm/step) to ensure adequate sampling. The tunable laser was set to 2% and the 1045nm line was set to 13%. SHG images were analyzed using the OrientationJ plugin (http://bigwww.epfl.ch/demo/orientation/). Briefly, OrientationJ Analysis was used with a local window σ of 1 pixel with a Fourier gradient to color code the images (Color survey HSB: Hue (Orientation), Saturation (Coherency), Brightness (Original-Image). OrientationJ Distribution was used to generate the distribution histogram and table of the orientation values (degrees). Cumulative distributions were compared using an unpaired two-tailed Kolmogorov-Smirnov test with 95% confidence level using the “Quest Graph Kolmogorov-Smirnov (K-S) Test Calculator” (AAT Bioquest, Inc., 26 Apr. 2023, https://www.aatbio.com/tools/kolmogorov-smirnov-k-s-test-calculator).

### Decellularization process and decellularized ECM (dECM) coating

Primary tumors were harvested, and viable tumor tissue was isolated and snap frozen in liquid phase nitrogen. Frozen tumor pieces were pulverized with cryogenic tissue crusher (Biospec Products) and hammer, on dry ice. Tissue powder was resuspended in 20mM EDTA (Quality Biological) with 2% Triton X (Fisher Scientific) in distilled water, at a concentration of 0.01- 0.05g/mL. Samples were adequately mixed on rotator overnight at 4C. Samples were spun down at 300xg for 5 minutes in 4C temperature-controlled centrifuge, and resulting pellet was resuspended in 1X PBS. Sample was washed with 1X PBS three more times. After the final wash, pellet was resuspended in 1mL of 1X PBS with 10X penicillin-streptomycin overnight at 4C. Samples were spun down at 300xg for 5 minutes in 4C temperature-controlled centrifuge, and resulting pellet was resuspended in 1X PBS, repeated three times. After the final wash, pellet was resuspended in 1X PBS. Samples were sonicated at 10amps for 10 seconds, three times. Total protein concentration was quantified by Bicinchoninic acid assay (BCA, Thermo Fisher) and prepared at 0.5-1.0mg/mL in 1X Dulbecco’s PBS (DPBS, Cytiva Hyclone) and then plated into 96-well tissue-culture treated plates (CytoOne) for dECM coating overnight at 37C, 5% CO2. Each biological dECM sample was plated in technical triplicates for the co-culture, and another set of technical triplicates to assess how much dECM coated the plate (by BCA). For the co-culture wells, 35,000 murine breast cancer cells were plated per well in 150uL complete media. The murine breast cancer cell line used for co-culture corresponded to orthotopic tumor in which dECM was derived from. For the wells used to assess how much ECM was coated, washed with 1X DPBS and aspirated all liquid left to dry, then ran BCA on these wells. Co-culture wells were incubated for 24 hours in 37C, 5% CO2.

### RNA Extraction and qRT-PCR Analysis

Tumor cell/dECM co-cultures were washed with DPBS and lysed inside the wells using Trizol (Invitrogen). RNA was isolated using Trizol as per manufacturer’s instructions (Chloroform, Sigma Aldrich). RNA was purified using Zymo RNA Clean & Concentration RNA Extraction Kit (ZYMO), as per manufacturer’s instructions. RNA quality and quantity by analyzed using a NanoDrop instrument, and RNA concentration was equalized for all samples prior to reverse transcriptase reaction. Reverse transcriptase reaction was performed using Bio-Rad iScript cDNA Synthesis kit (Bio-Rad), following manufacturer’s instructions. Bio-Rad SYBR Green qPCR (Bio-Rad) was performed in hard-shell 384-well PCR plates (Bio-Rad), run in triplicate to assess changes to gene expression of Snail, Twist, Slug and Zeb EMT transcription factors, and beta-actin housekeeping gene. Bio-Rad CFX Manager 3.1 software was used on Bio-Rad CFX96 Touch Real-Time PCR Detection System. Fold-change was calculated by ΔΔCT Livak’s method. Samples that lacked high RNA purity standards were excluded from the analysis (< 1.8 OD).

Primer sequences (IDT):

Snail

Fwd aagatgcacatccgaagc

Rev atctcttcacatccgagtgg

Twist

Fwd agctacgccttctccgtct

Rev tccttctctggaaacaatgaca

Slug

Fwd ctccagaccctggctgcttca

Rev gtctgcagatgtgccctcagg

Zeb

Fwd tgagcacacaggtaagaggcc

Rev ggcttttccccagagtgca

Beta actin

Fwd ctaaggccaaccgtgaaaag

Rev accagaggcatacagggaca

### Western Blot of EMT transcription factors

The cell lysates were collected using RIPA buffer supplemented with protease inhibitors (Roche, 11697498001). Briefly, proteins were separated by 12% SDS-PAGE electrophoresis and then transferred to PVDF membranes (GE Healthcare). After blocking with 5% BSA for 1 hour, the membranes were incubated with primary antibodies against Actin (Millipore Sigma, MAB1501, 1:1,000), Twist (Proteintech, 25465-1-AP, 1:500), Zeb1 (Santa Cruz Biotechnology, SC-25388, 1:500), Snail (Cell Signaling Technology, 3879S, 1:500), Slug (Cell Signaling Technology, 9585S, 1:500) overnight at 4°C.

After washing three times, the blots were further incubated with donkey anti-rabbit IRDye 800CW (LI-COR, 925-32213, 1:5000) or donkey anti-mouse IRDye 680RD (LI-COR, 925-68072, 1:10000 secondary antibody accordingly. The images were detected by an Odyssey infrared imaging scanner with Image Studio V5.2 software (LI-COR, USA).

### 3D Hydrogel Model

Untreated primary mammary tumors were harvested, and tumor cell clusters isolated using low concentrations of collagenase and trypsin (Life Technologies). Isolated tumor cell clusters were cultured in low adhesion plates (Corning) for 24 hours (37C, standard cell culture conditions) before using in static 3D hydrogels or 3D microfluidic model systems. Tumors from control and Treg cell ablated mice (MMTV-PyMT, PyMT and EO771) were minced with a scalpel and then agitated in solutions of 0.1% Triton (overnight, Fisher Scientific), 2% sodium deoxycholate (overnight, Sigma Aldrich), sodium chloride (1 hr, FUJIFILM Wako Pure Chemical Corp), and DNAse (overnight, Sigma Aldrich) ^22^. The decellularized tumor matrix scaffolds (dECM) were frozen in -80C overnight and then lyophilized. Lyophilized tissue was minced with a scalpel into pieces about 1mm in diameter and stored at -20C until pre-gel formation. Pre-gel was formed by adding 1% decellularized tissue to 0.01M HCl with 0.1% pepsin (Sigma Aldrich). Pre-gel was mixed for 24 hr, chilled, neutralized with 10X PBS and 0.1M NaOH (Sigma Aldrich), and used immediately in static 3D hydrogels and 3D microfluidic models as described ^23^. Briefly, the pre-gel dECM was mixed with fibrin (20 mg/ml fibrinogen with 3 U/ml thrombin, Sigma Aldrich) at a 1:1 ratio, and mixed with PyMT or EO771 primary tumor clusters, respectively. This mixture was cultured in a PDMS (polydimethyl-siloxane, Fisher Scientific) ring attached to a glass coverslip in a 24-well plate and allowed to polymerize at 37C for 30 minutes before adding cell DMEM culture media (Thermo Fisher) supplemented with 10% FBS and Pen/Strep antibiotics (Fisher Scientific). Samples were incubated at 37C under standard cell culture conditions until invasive, about 2 days. For proliferation evaluation, tissues were cultured for 24 hr or 48 hr, fixed in 4% paraformaldehyde and blocked in 2% BSA in PBS with 0.1% Tween20. Tissues were stained for phospho-histone H3 (PH3, Ser10) antibody (Invitrogen PA5-17869, 1:500x) and Alexa Fluor 488 phalloidin (Thermo Fisher A12379, 1:500x) overnight (4C). Secondary antibody Goat Anti-rabbit Alexa Fluor 568 (1:500x, 2hr incubation at 4°C). DAPI counterstaining was used for nuclei. A minimum of 3 biological replicates were performed for each experiment, each with minimum of 3 technical replicates.

3D static hydrogels were imaged using Zeiss LSM980 confocal microscopy with ZEN 3.3 software at 20x magnification, with slices taken every 10μm. Orientation of collagen fibers were analyzed using ImageJ OrientationJ plugin and custom MATLAB script (Taufalele et al, 2019). Tumor cell cluster area and perimeter were calculated in ImageJ using the “Analyze Particles Shape Descriptors Function”. Z-projections of DAPI and the PH3 were used to quantify the area of DAPI and PH3 signal using FIJI/ImageJ Analyze particles shape descriptors. The %PH3 per spheroid was calculated by dividing the PH3 area by the DAPI area per spheroid.

### 3D Microfluidic Model System

The same pre-gel dECM with fibrin and cells mixture used in static 3D hydrogels was loaded into our 3D microfluidic model system ^23^, allowed to polymerize at 37C for 30 minutes before adding cell culture media (DMEM+10%FBS+P/S). Devices were cultured in low oxygen conditions (5% O2) for 48 hours before inducing biochemical gradient of stromal derived factor-1 (SDF-1, Sigma-Aldrich) established to promote collective migration, as previously described ^23^. Live-cell imaging was performed using a Nikon Ti-E microscope with NIS-Elements software at 37C, and 5% oxygen, at a 63x magnification. Cell clusters were marked using Nikon Imaging software and pictures were taken every 20 minutes for a maximum of 12 hours. We analyzed 15-20 organoids from at least 3 different mice (4-8 organoids per mouse) to perform statistical analysis (determined via power analysis with 50% change between experimental and control groups, a 0.05 significance level and 0.80 power). Image analysis was performed using Nikon Imaging Software and MATLAB to generate cluster migration maps and quantify migration efficiency in the direction of the gradient, and average velocity (uM/min) ^23^.

### Flow Cytometry Analysis

#### Analysis of Circulating Tumor Cells (CTCs)

Whole blood was collected at the indicated end points via cardiac puncture on isoflurane-anesthetized mice, followed by euthanasia. Blood was lysed in ACK red blood cell lysis buffer (150mM NH4Cl, Fisher Chemical; 10mM KHCO3, Fisher Chemical; 0.1mM EDTA, Quality Biological pH 7), and samples were spun down at 350xg, for 10 minutes in 4C temperature-controlled centrifuge. The resulting supernatant was aspirated, and cell pellet was resuspended in 2mL of ACK red blood cell lysis buffer. Samples were spun down at 350xg, for 10 minutes in 4C temperature-controlled centrifuge. The resulting supernatant was aspirated, and the cell pellet was resuspended in 2mL of 1X PBS. Samples were spun down in 1300 RPM for 8 minutes. The resulting supernatant was aspirated, and cell pellet was resuspended in 300uL of MACS buffer (0.5% BSA, 2mM EDTA in 1X PBS, pH 7).

Samples were transferred into a 96-well V-bottom plate (USA Scientific) and incubated with 50uL of Zombie Aqua viability dye (BioLegend Cat#432101, 1:1000) in PBS buffer for 15 minutes, covered on ice. Plate was spun down at 1350RPM for 7 minutes, in 4C temperature-controlled centrifuge. The resulting supernatant was dumped, and pellets were resuspended in 50uL of Fc block (TONBO Bioscience Cat#70-0161-U500, 1:10) in MACS buffer for 10 minutes, covered on ice. Samples were stained with AF-700-conjugated anti-CD45 antibody (30-F11 clone, Thermo 56- 0451-82, 1:125) in MACS buffer for 20 minutes, covered on ice. Wells were washed with MACS buffer and resuspended in 100uL of fixation/permeabilization solution (TONBO, TNB-0607-KIT) for 40 minutes, covered on ice. Cells were then permeabilized with 1X Perm Buffer (TONBO, TNB- 0607-KIT), for 20 minutes, covered on ice. The plate was spun down at 1550 RPM for 7 minutes, in 4C temperature-controlled centrifuge. Cell pellets were resuspended in 50uL of anti-cytokeratin PE (C-11 clone, Thermo Fisher MA5-28574, 1:100 in 1X Perm Buffer) for 30 minutes, covered on ice. Cells were washed and were resuspended in 300uL of 1X PBS, filtered through 35uM mesh (Fisher), and 10uL of Counting Beads (BioLegend, Cat#424902) were added to each sample for analysis. Samples were run in a LSR Fortessa (BD) with FACS Diva software at the VCU Flow Cytometry Core. Analysis was performed using Flowjo software (Tree Star). Compensation and positive and negative controls were prepared using lysed whole blood from tumor-naïve mice spiked-in with known amounts of murine breast cancer cell lines, or no added cells.

#### Analysis of Immune Cell Identification by LPS and DT Treatment

Blood and spleen samples were collected from mice for DT treatment verification and LPS stimulation experiments, respectively. Blood samples were processed as described above to generate single-cell suspensions. Spleen samples were crushed between frosted glass slides with 5mL 1X PBS, to generate single-cell suspensions. Samples were filtered through a 100 µm strainer, then centrifuged at 400g for 5 minutes. The resulting supernatant was aspirated, and cell pellet was resuspended in 2mL of ACK red blood cell lysis buffer, incubated for 5 minutes on ice. Samples were spun down at 350xg, for 5 minutes in 4C temperature-controlled centrifuge. The resulting supernatant was aspirated, and the cell pellet was resuspended in 100uL of MACS buffer. Samples were stained with viability dye and cell surface markers as described above. For viability, we used Zombie NIR viability dye (BioLegend Cat#423106, 1:1000) in PBS buffer. The following antibodies were used: anti-CD45 BUV395 (30-F11 clone, BD Biosciences 564279, 1:200), anti-CD11b PE/Fire640 (M1/70 clone, BioLegend 101280, 1:200), anti-Ly6C PE (HK1.4 clone, BioLegend 128008, 1:250), anti-Ly6G BV605 (IA8 clone, BioLegend 127639, 1:200), anti-F4/80 BUV496 (T45-2342 clone, BD Biosciences 750644, 1:200), anti-CD11c BV711 (N418 clone, BioLegend 117349, 1:200), anti-TCRb AF700 (H57-597 clone, BioLegend 109224, 1:200), anti-CD4 BUV737 (GK1.5 clone, BD Biosciences 612761, 1:200), anti-CD8 APC/Fire750 (53-6.7 clone, BioLegend 100766, 1:200), anti-CD335 PE/Cy7 (29A1.4 clone, eBiosciences 25-3351-82, 1:200), anti-CD25 BV421 (PC61 clone, BD Biosciences 562606, 1:200), anti-CD19 BV650 (6D5 clone, BioLegend 115541, 1:200), anti-TCRgd BV786 (GL3 clone, BD Biosciences 744117, 1:200), anti-CD69 PE (H1.2F3 clone, eBiosciences 12-0691-81, 1:200); all antibodies were diluted in 1X MACS buffer. Instead of fixation/permeabilization, cells were resuspended in 100uL of 1X PBS. Samples were run in a Cytek Aurora with SpectroFlow (Cytek) software at the VCU Flow Cytometry Core. Analysis was performed using Flowjo software (Tree Star).

### Collagenase Activity

The biological activity of the expressed L. sericata collagenase (MMP-1) was assessed using Collagen Degradation/Zymography Assay Kit (ab234624, Abcam, USA) according to the manufacturer’s instruction. In brief, 0.02g of fresh tumor tissue was subjected to lysis in 200μL of cell lysis buffer. Following purification, the lysate was activated using 1mM APMA (ab112146, Abcam, USA) at 37℃ for 3 hours. Subsequently, 50μL of the activated lysate was dispensed into the designated well(s) of a 96-well plate (655098, Greiner Bio-One, Germany), and the volume was adjusted to 100μL by adding 50μL of collagenase substrate. The plate was then inserted into the POLARstar OPTIMA microplate reader, and the fluorescence of the assay was recorded every minute at Ex/Em 490/520nm in kinetic mode at 37℃ for 2 hours. Finally, the activity of the sample was determined using the calculation formula provided by the kit.

### Bioluminescence imaging

Lung metastatic burden was determined by bioluminescence imaging using an IVIS-200 imaging system (Xenogen Corp.) with Living Image 4.7.4 software. To this end, mice were anesthetized and injected retro-orbitally with 1.5mg of D-Luciferin (15mg/ml, Syd Labs) and imaged 2 minutes after injection. For ex vivo lung metastatic burden determination, mice were euthanized 2 minutes after D-Luciferin injection, and lung imaging completed within the next 5 minutes.

### RNA-Seq Analysis

FastQC v.0.11.5 was used for quality control at all stages of RNA-seq preprocessing. The Mouse GRCm38/mm10 reference genome was obtained from UCSC Genome Browser Gateway (http://hgdownload.soe.ucsc.edu/goldenPath/mm10/bigZips/chromFa.tar.gz). The gene annotation file was obtained from Gencode (https://ftp.ebi.ac.uk/pub/databases/gencode/Gencode_mouse/release_M25/gencode.vM25.annot <u>ation.gtf.gz</u>) on 09/03/2022. Only autosomes, mitochondrial, and sex chromosomes were used. Reads were trimmed with Trim Galore! v0.6.4_dev and aligned to the indexed reference genome using bwa mem v0.7.17-r1188 (Li and Durbin, 2009). Read counting on a transcript reference file was performed using featureCounts v2.0.1 (Liao *et al.*, 2014). The counts were imported into R v.4.2.1 and the log2 transformed Transcripts per Million (TPM) values were calculated using an in-house script.

A differential expression analysis was performed on control or Treg cell-ablated EO771 primary tumors using raw reads counts. Genes with a read count of zero for all samples were removed prior to identifying differentially expressed genes using the edgeR R package v3.40.2 (Robinson *et al.*, 2010). Genes are considered significantly differentially expressed if the False Discovery Rate adjusted p-value is < 0.05.

### Matrisome Gene Signature, Average Profile, and Human Orthologs

A list of genes that encode ECM and ECM-modifying proteins was compiled from the literature ^28-31^. A signature was identified by intersecting this list with the list of significantly differentially expressed genes (FDR <=0.05 and log2 FC >=1.4 or <=-1.4) from the control and DT-treated bulk RNASeq data from primary tumors. The HGNC Comparison of Orthology Predictions (HCOP) ^52,53^ was used to identify human orthologs by mapping the mouse Ensembl gene identifiers to human Ensembl gene identifiers and official gene symbols. HCOP was set to identify orthologs between Mouse as the source and Human as the ortholog with all default ortholog databases being utilized, which identified 83 human orthologos for the 73 matrisome mouse genes. Data was downloaded as a text file. All list operations and the average matrisome signature profile calculation was performed in R v.4.2.1.

### TCGA Gene Expression and Survival Analysis

Gene expression and survival data for human cases in The Cancer Genome Atlas (TCGA) ^54^ were obtained utilizing the methods described previously ^55^. Briefly, legacy RSEM scaled estimates for the breast cancer (BRCA) TCGA cohort were downloaded using the Bioconductor R package TCGABiolinks v.2.5.9 ^56^. Scaled estimates were converted into log2 transcripts per million (TPM) values and quartile normalized. This TPM matrix was then filtered to include only the 46 human orthologs of the 40-gene mouse matrisome signature and row-median centered. The ComplexHeatmap v2.16.0 ^57^. R package was utilized for K-means clustering by setting the “column_km” parameter to 3, and for final heatmap visualization. The TCGA-BRCA cohort contains survival data for 1078 cases. Those with data beyond 10 years (3650 days) were censored as “alive” at the 10-year mark. The survival v3.5-5 ^58^ and survminer v0.4.9 (https://rdrr.io/cran/survminer/.) packages were used to generate Kaplan-Meier survival curves, Log-Rank p-values, and hazard ratios.

### Single-cell RNA-seq Analysis on Primary Tumors

#### Tumor sample preparation for scRNA-seq

The primary mammary gland tumor was finely diced and minced until it reached a paste-like consistency. Subsequently, the minced tumor was transferred into a 15ml tube and underwent digestion for 30 minutes at 37°C using Liberase™ TL (Roche). Following digestion, the tissue was suspended in RPMI supplemented with 10% fetal bovine serum (FBS) and filtered through a 100 µm strainer, then centrifuged at 400g for 10 minutes. The resulting pellet was resuspended in 10ml of 43% Percoll™ Centrifugation Media (Cytiva) and centrifuged at 500g for 10 minutes. The supernatant, containing fat and other impurities, was carefully removed, and the cell pellet was washed with 10ml of PBS. Finally, the cells were treated with ACK buffer to remove any remaining red blood cells. The total and viable single cell numbers were counted, with samples boasting cell survival rates of over 80% being chosen for further procedures. The selected cells were washed and suspended in PBS, reaching a final concentration between 700 and 1200 cells/μL, adhering to the protocol outlined in the Chromium Next GEM Single Cell 3’ V3.1 User Guide for operation on the 10x Genomics Chromium system. Gel Beads-in-emulsion (GEMs) were then formed to isolate single cells. After successful GEM formation, reverse transcription was carried out in a PCR machine to enable labeling. Subsequently, magnetic beads were utilized to purify and enrich the first strand cDNA. The purified cDNA underwent amplification and quality control steps. Using the 10x Genomics library Construction Kit, libraries were constructed from quality-verified cDNAs by following the User Guide. Finally, the library was submitted for sequencing.

#### Custom Mouse/Firefly Luciferase Reference Genome for Single Cell Alignment

To identify tumor cell clusters in the single cell data we created a custom mouse reference genome that included the Firefly Luciferase gene sequence. These tumor cells happen to express this non-mouse gene, and are thus, easily identified by this feature. Briefly, the P.Pyralis Luciferase gene sequence with accession M15077.1 was downloaded from GenBank on January 31, 2022 (https://www.ncbi.nlm.nih.gov/nuccore/M15077.1). This sequence was appended to the end of the fasta and GTF files for the 10X Genomics CellRanger mm10-2020-A version of the mouse genome that is from GENCODE version M23 (https://www.10xgenomics.com/support/software/cell-ranger/downloads/cr-ref-build-steps#mouse-ref-2020-a) and then run through the reindexing process as described by 10X Genomics documentation. This custom merged reference genome was utilized for alignment of all single cell samples.

#### Single Cell RNA-Seq Analysis

Single cell samples were processed and analyzed utilizing a combination of community tools, 10X Genomics software and in-house R scripts for quality control, alignment, and downstream analyses. Briefly, sample-level QC was performed on the raw FastQ files using FastQC v0.11.9 (Andrews 2018) and MultiQC v1.11 (Ewels et al. 2016). The 10X Genomics CellRanger v6.0.1 software was used for alignment to a custom mouse/firefly genome. Poor quality and dying cells were identified with a custom R script that utilized R v4.1.3 and the Seurat v4.3.0 R package. Metrics utilized include mitochondrial gene expression percentage, number of detected genes, and the count of unique molecular indices (UMIs). For each sample, any cell above or below 3 median standard deviations (MADs) for the number of genes or UMI counts were considered poor quality or extreme outliers and excluded from analysis. Additionally, any cells above 3 MADs for the mitochondrial expression were considered to be dying and removed from the dataset. After identification and removal of poor-quality cell barcodes, samples were normalized and merged using Seruat’s “merge()” function with log normalization and scaling in a custom R script. This script also generated tSNE and UMAP layouts, performed a cell cycle analysis, clustering, and utilized the LoupeR v1.1.0 package to convert the processed data into a Loupe file compatible with 10X Genomics Loupe Cell Browser v7 (https://www.10xgenomics.com/support/software/loupe-browser/latest).

#### Single Cell RNA-Seq Cell Identification Analysis

We first ran global differential expression analysis between each cluster versus all the other clusters in the entire dataset on 10X Genomics Loupe Cell Browser v7. From this output, we sorted on Log2 FC by descending order, for each respective cluster. For each cluster, we collected the top differentially expressed genes (DEGs) that were unique to that particular cluster. Unique was defined as: cluster of interest global Log2 FC > or = to 2 times global Log2 FC for other clusters. Next, the top DEGs for each cluster were input into the Mouse Molecular Signatures Database (MSigDB) on the Gene Set Enrichment Analysis (GSEA) web platform, to compute overlaps between our top DEGs with other publicly available datasets deposited to GSEA. We computed overlaps between our top DEGs and other datasets denoted as cell type signature gene sets, with FDR q-value less than 0.05 for top 50 gene sets. The output showed what cell type signature gene sets had significant overlap with our top DEGs, which hinted towards cell identities for each cluster. To further validate the identity of each cluster, we overlaid identity marker genes on cell clusters using 10X Genomics Loupe Cell Browser v7 (Fig.S8).

#### Single Cell ECM Signature Scoring

A custom in-house R script and algorithm was developed to calculate a single score per cell to reflect that cell’s expression of the genes in the ECM gene signature identified using the bulk RNASeq data. First the ECM signature gene count data was normalized using Seurat’s ScaleData function with options to utilize the Negative Binomial distribution, turn centering off, and regress out the UMI count to adjust for the library size of each cell as tumor and fibroblast cells tended to have a much higher UMI than other cells. Signature scores were then calculated as the sum of the scaled expression value across all genes in the signature. A cell was labeled as having the signature if the score was greater than or equal to the 3rd quartile, with the quartiles being calculated across all cells in the dataset. Cells with scores less than the 3rd quartile were labeled as not reflecting the signature. The signature scores and categories were overlaid onto the UMAP plots using Seurat’s featurePlot() function. The categorical data was also uploaded into the Loupe Cell Browser for additional visualization.

### Statistical Analysis

All statistical analysis was performed using GraphPad Prism V10 software. Unpaired and paired Student’s t-tests, one-way analysis of variance (ANOVA) with post-test Tukey HSD’s multiple comparisons, and two-way ANOVA with multiple comparisons were used as described in the legend figures. Non-parametric analyses were performed when appropriate and denoted in the figure legend of the corresponding figures. Results are presented as mean +/- SEM for parametric data, or median +/- interquartile range for non-parametric data. p-values of less than 0.05 were considered statistically significant. ***

## DATA AVAILABILITY

The bulk RNA-seq datasets generated during the current study is available on GEO (GSE240408, reviewer’s access token mtytcmkcnvadbkh). The single-cell RNA-seq dataset GEO submission is in process.

## Supplement Figure Legends

**Video S1: Representative video of a PyMT tumor organoid seeded in control PyMT dECM responding to an SDF-1 gradient on a 3D microfluidic system.**

**Video S2: Representative video of a E0771 tumor organoid seeded in control E0771 dECM responding to an SDF-1 gradient on a 3D microfluidic system.**

**Video S3: Representative video of a PyMT tumor organoid seeded in Treg cell ablated PyMT dECM responding to an SDF-1 gradient on a 3D microfluidic system.**

**Video S4: Representative video of a E0771 tumor organoid seeded in Treg cell ablated E0771 dECM responding to an SDF-1 gradient on a 3D microfluidic system.**

**Figure S1. DT treatment to Foxp3DTR mice ablates Treg cells, with no changes to other immune cell populations.** (A) Schematic of experimental design. Black downward arrow indicates orthotopic implantation, green arrows indicate DT treatment, black upward arrow indicates blood collection. (B) Gating strategy for flow cytometry panel to assess changes to immune cell populations following DT treatment. Dictionary table for cell identities and corresponding markers. (C) Contour plot of Foxp3/CD25 expression on CD4 T-cells, for representative control and DT treated blood samples. Quantification of percentage (%) Treg cells from CD4 T-cell parent population. Data is displayed as mean +/- SEM. *p<0.05. (D) Quantification of immune cell % for control (black) and DT treated (green) samples. % of CD45 cells, CD4 T-cells, CD8 T-cells, gamma delta T-cells, NK cells, B-cells, DCs, monocytes, granulocytes, and macrophages are shown. Respective parent populations are noted on the y-axis title. Data is displayed as mean +/- SEM. n = 2 mice per condition.

**Figure S2. LPS model validation by immune cell activation markers in flow cytometry. LPS-stimulated animals show an increase in Treg cell populations, and enrichment of collagen amounts in orthotopic PyMT tumors.** (A) Schematic of experimental design. Black downward arrow indicates orthotopic implantation, green arrows indicate LPS treatment, black upward arrow indicates tissue collection. (B) Gating strategy for flow cytometry panel to confirm systemic inflammation following LPS treatment, by assessing activation markers CD25 and CD69 on B-cells, macrophages, and T-cells. (C) Histograms of CD69 expression on CD4 T-cells, CD8 T-cells, B-cells, and macrophages; and CD25 expression on CD4 T-cells and macrophages. Representative control (grey) and LPS treated (lavender) spleen samples overlaid. Quantification of percentage (%) of CD4 T-cells that express CD69, % of CD8 T-cells that express CD69, % of B-cells that express CD69, and median fluorescence intensity (MFI) of CD69 on macrophages. Quantification of % of CD4 T-cells that express CD25, and MFI of CD25 on macrophages. Data is displayed as mean +/- SEM. (D) Contour plot of Foxp3/CD25 expression on CD4 T-cells, for representative control and LPS treated spleen samples. Quantification of % of Treg cells from CD4 T-cell parent population. (E) Representative images of Picrosirius Red staining of primary orthotopic PyMT tumors for control and LPS treated conditions. Scale bar = 250 um. Quantification of collagen-positive area in PyMT orthotopic tumors. n = 2-4 mice per condition. Analyzed one bilateral orthotopic tumor, per mouse. 10 representative images at 20x magnification were analyzed per mouse using a combination of ImageJ. Data is displayed as mean +/- SEM. **p<0.01. Statistical analysis was performed by unpaired t-test with Welch’s correction. Less collagen amounts were observed upon Treg cell-ablation (Fig 2C), whereas systemically induced inflammation via LPS treatment led to more collagen amounts compared to control tumors. Interestingly, we also show that LPS-stimulation led to an increase in %Treg cells.

**Figure S3. No changes to proliferation in spheroid cells cultured with dECM for static 3D hydrogel experiments.** (A) Quantification of percentage (%) of cells that express phospho-histone 3 (PH3) in spheroids used for static 3D hydrogel experiments. Spheroids that were cultured in control dECM are in black, and spheroids cultured in Treg cell-ablated dECM are in green, for both E0771 and PyMT cell lines. Signals were collected at 0- and 48-hour time-points, and analyzed with ImageJ using particle shape descriptors. (B) Quantification for tumor cluster perimeter and (C) area, at 0- and 48-hour time-points. Data is displayed as mean +/- SEM. * p<0.05, **p<0.01, ***p<0.001. Statistical analysis was performed by Mann-Whitney tests.

**Figure S4. Effective and reproducible decellularized ECM (dECM) coating onto TC-treated wells.** (A) Representative images of coated wells with control tumor dECM (black), DT-treated tumor dECM (green), or an empty well (pink). Imaged on Keyence brightfield, at 20X magnification. (B) Representative images of DAPI-stained plated cells (top) and coated dECM well (bottom). (C) Schematic showing that after dECM samples were normalized to 0.5-1mg/mL, they were plated in triplicates for assessment of how much dECM was coated onto the plate by BCA total protein quantification. (D) BCA standard curve points (black) with linear regression curve (black line), run on TC-treated plate in which dECM samples were coated on. R-square and regression equation are shown on the graph. Inter-polated dECM samples (red) that coated onto TC-treated plates are plotted on-top of the BCA linear curve. Dotted line represents the calculated concentration for empty wells (on TC-treated plate) washed with 1X PBS, denoted as “Bkgd Conc.” Example calculation for determining how much dECM was coated on the TC-treated plate is shown for Control2. Data shown is for PyMT Plate #2, which is representative of data for the other plates. (E) Percent coefficient of variation (%CV) for OD values of coated dECM samples (as determined by BCA), across technical replicate wells. Data is displayed as mean +/- SEM. Data shown is for all plates and standard points. This panel shows that dECM coating onto TC-treated plates had reasonable variation across technical replicates, for the most part < 10%CV.

**Figure S5. EMT transcription factor western blot in PyMT and E0771 cell lines.** Western blotting of Zeb (≥180 kDa corresponding to the post-translational modified forms & 124 kDa unmodified form), Twist (25-30 kDa), Snail (29 kDa), and Slug (30 kDa) EMT transcription factors, as well as B-actin (42 kDa) as a house-keeping control. With exception of Slug in PyMT cells, all other transcriptions factors are expressed in both cell lines.

**Figure S6. Validation and gating strategy of CTC flow cytometry panel on tumor-naïve whole blood, and tumor cell-spike in.** (A) Schematic of gating strategy for CTC panel on Unstained Tube (Tumor Naïve Whole Blood). Gated on cells, singlets, live cells, CD45-, and Cytokeratin+ cells. We show that excluding low FSC/SSC particles results in no Cytokeratin positive population, as expected. (B) Same as A, for Stained Tube (Tumor Naïve Whole Blood). We show bright, distinct CD45+ population, as expected from whole blood samples. Additionally, we show no Cytokeratin positive population, as expected from a tumor-naïve animal. (C) Same as A, for Stained Tube (Tumor Naïve Whole Blood + Tumor Cell Spike In). We show bright, distinct CD45+ population, as expected from whole blood samples. Additionally, we show bright, distinct Cytokeratin+ population, as expected from spiking-in Tumor Cells.

**Figure S7. Collagenase activity from PyMT and E0771 orthotopic tumors.** (A) Representative collagenase kinetic curve from PyMT control (black) and Treg cell ablated (green) tumors. Curves show more collagenase activity among the Treg cell ablated tumors, compared to Control tumors. (B) Quantification of collagenase activity in E0771 (left) and PyMT (right) orthotopic tumors. n = 4 mice per condition. Analyzed one bilateral orthotopic tumor, per mouse. Technical replicates are shown, data is displayed as mean +/- SEM. (C) Quantification of collagenase activity in PyMT orthotopic tumors. Samples are denoted as Control (black), Treg cell ablated (green), IFN-γ neutralizing antibody (red), IFN-γ neutralizing antibody + Treg cell ablated (blue), *IFN-*_γ_*KO* Treg cell ablated (orange). n = 4-12 mice per condition. Analyzed one bilateral orthotopic tumor, per mouse. Technical replicates are shown, data is displayed as mean +/- SEM. Statistical analysis was performed by one-way analysis of variance with Tukey’s multiple comparisons post-test.

**Figure S8. PCA analysis shows successful grouping among biological replicates for control and Treg cell ablated samples. 5-year survival of clustered TCGA breast cancer patients show no statistically significant changes in overall survival.** (A) PCA graph showing clustering of E0771 samples (biological replicates shown) by overall gene expression using RNA-Seq. E0771 Control and Treg cell ablated samples were clustered together. (B) Kaplan-Meier survival analysis among clustered “Treg cell ablated-like” and “Control-like” TCGA breast cancer patients over a 5-year period using the 40-gene highly expressed matrisome signature. (C) Tables with patient survival information per cluster over time used for the Kaplan-Meier survival analysis censored at 5- and 10-year time points.

**Figure S9. Identity marker genes were used to identify scRNA-seq cell cluster identities.** (A) Bubble plot showing cell type-specific markers based on literature (y-axis), across different cell clusters (x-axis). Color gradient indicates the level of expression, and dot size indicates the percentage of cells that express the denoted marker. (B) UMAP colored by level of expression for each indicated marker, using 10X Genomics Loupe Cell Browser v7.

