## Supplementary material for "Tumor cell dissemination is facilitated through regulatory T cell-driven extracellular matrix remodeling": Fig. S1

Supplementary Figure 1

A

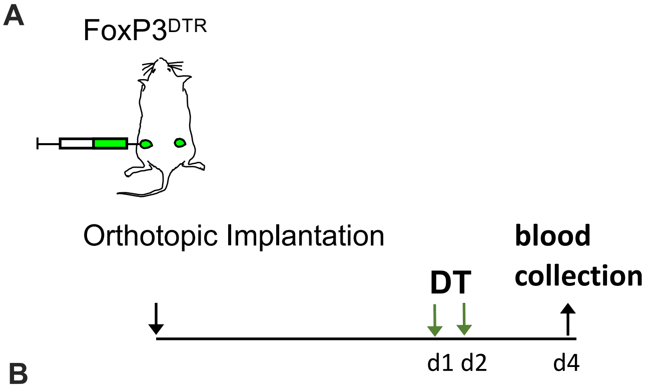


B

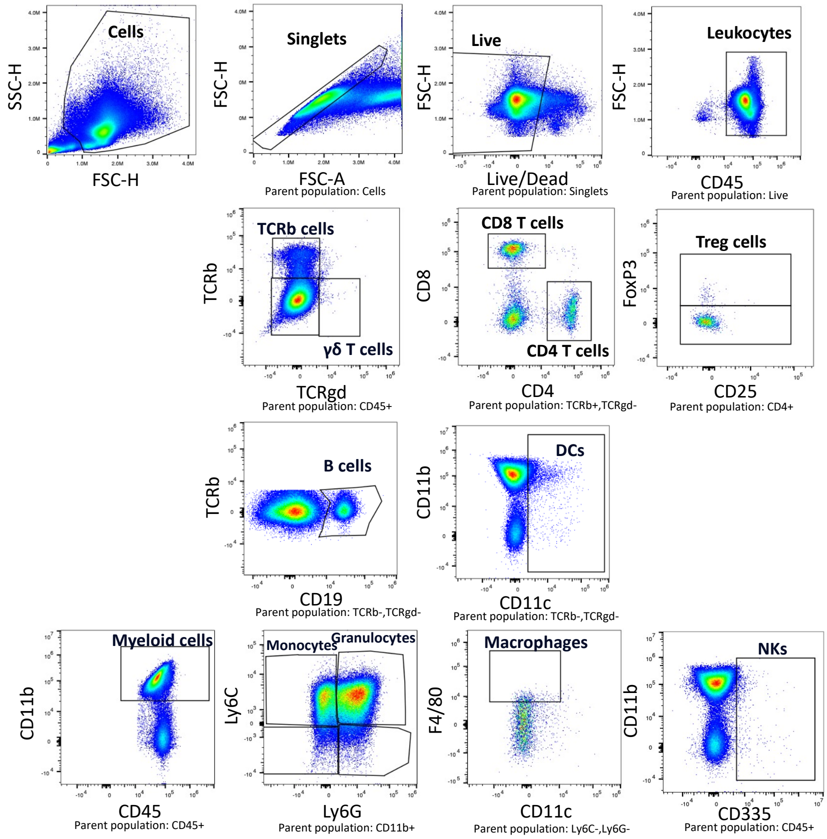


C

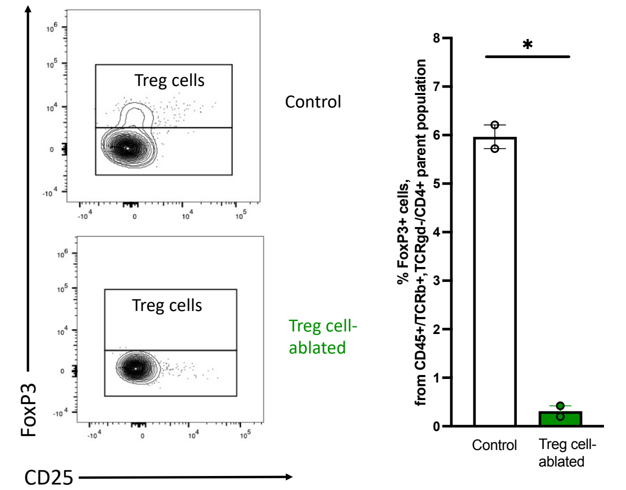


D

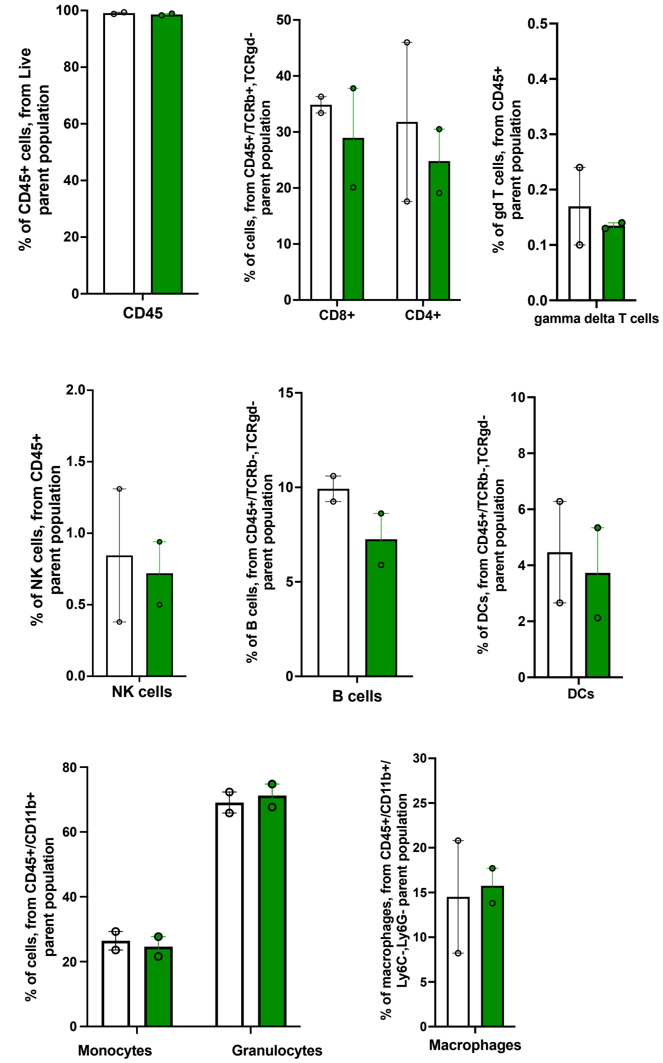


| Cell Identity | Markers |
|---|---|
| Cytotoxic T cells | CD45+/TCRb+,TCRgd-/CD8+ |
| Helper T cells | CD45+/TCRb+,TCRgd-/CD4+ |
| Treg cells | CD45+/TCRb+,TCRgd-/CD4+/FoxP3+ |
| Gamma delta T cells | CD45+/TCRb-,TCRgd+ |
| B cells | CD45+/TCRb-,TCRgd-/CD19+, MCHII+ |
| Dendritic cells | CD45+/TCRb-,TCRgd-/CD11c+ |
| Monocytes | CD45+/CD11b+/Ly6C+,Ly6G- |
| Granulocytes | CD45+/CD11b+/Ly6C+,Ly6G+ |
| Macrophages | CD45+/CD11b+/Ly6C-,Ly6G-/CD11c-,F4/80+ |
| NK cells | CD45+/CD335+,TCRb- |
| NK T cells | CD45+/CD335+,TCRb+ |
