## Supplementary figures and images for "Tumor cell dissemination is facilitated through regulatory T cell-driven extracellular matrix remodeling"

### Fig. S2

Supplementary Figure 2

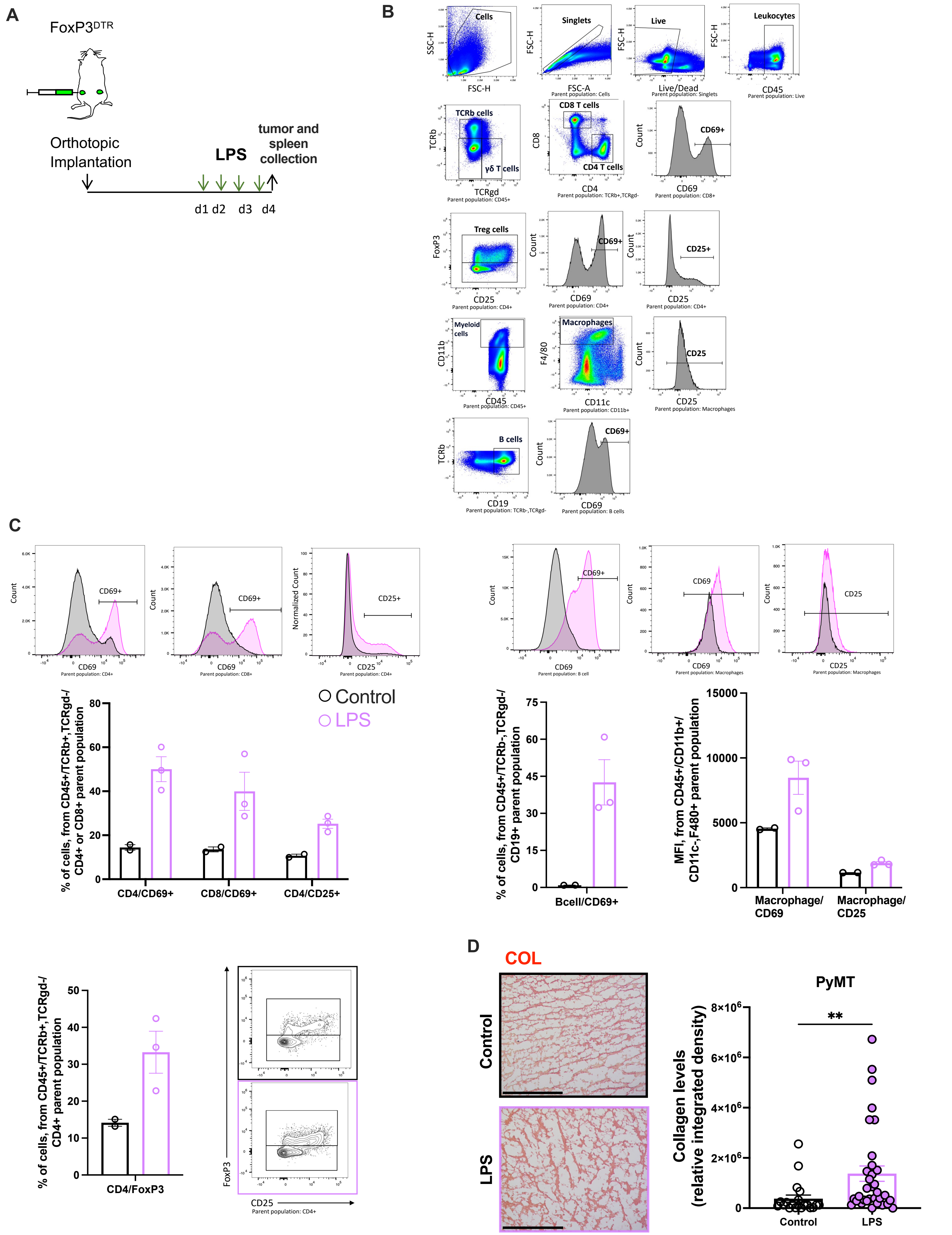

### Fig. S3

Supplementary Figure 3

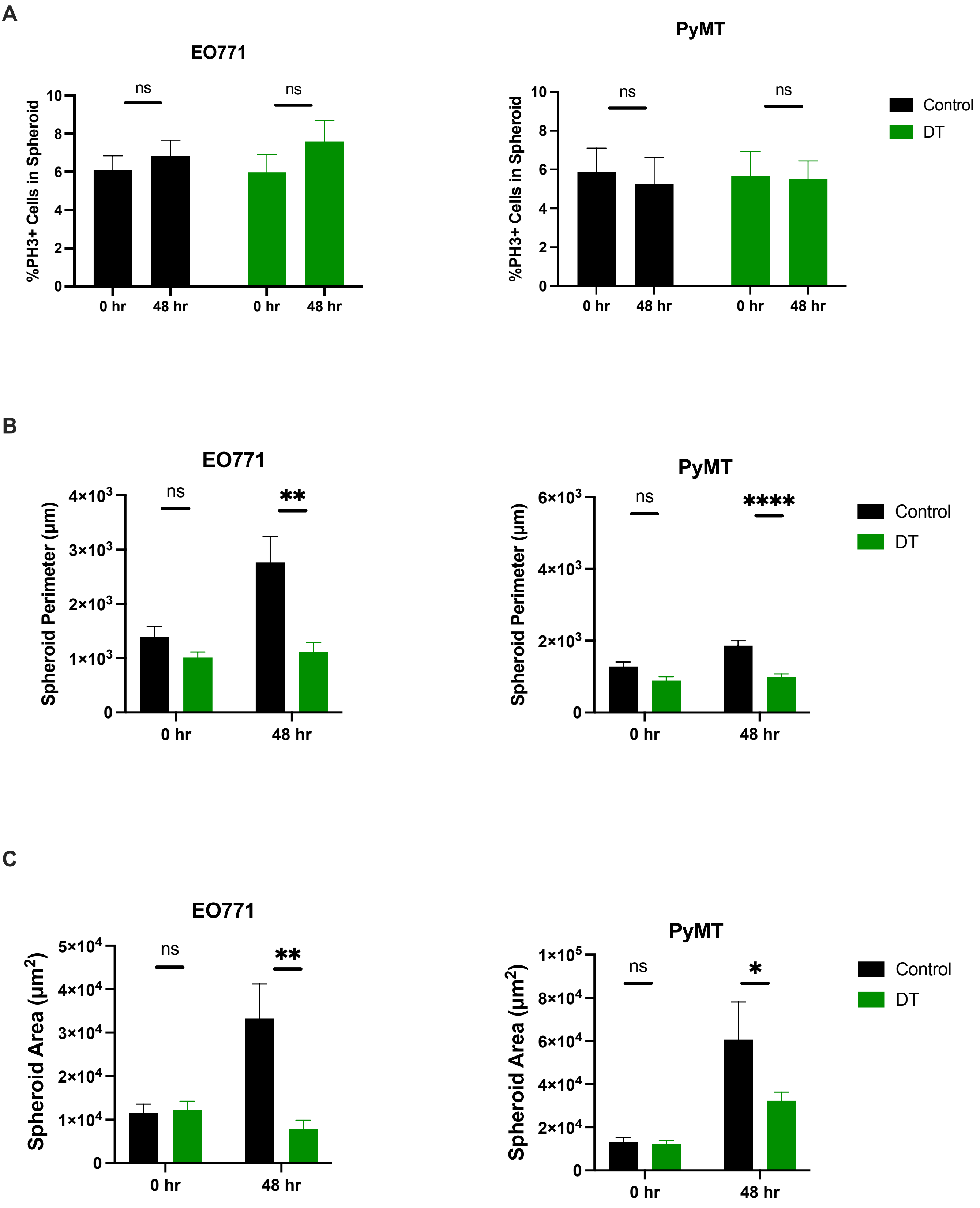

### Fig. S4

Supplementary Figure 4

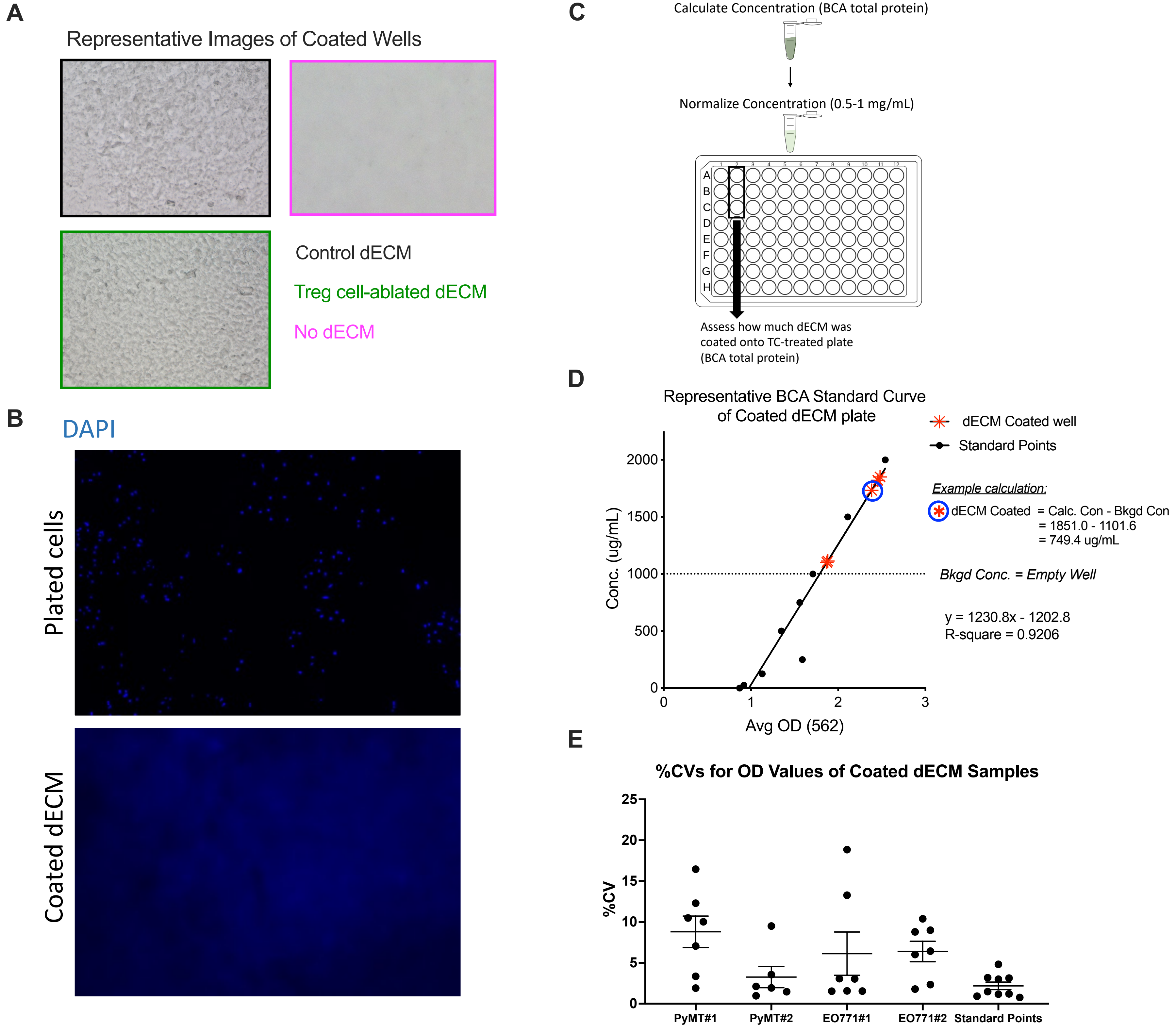

### Fig. S5

#### Supplementary Figure 5

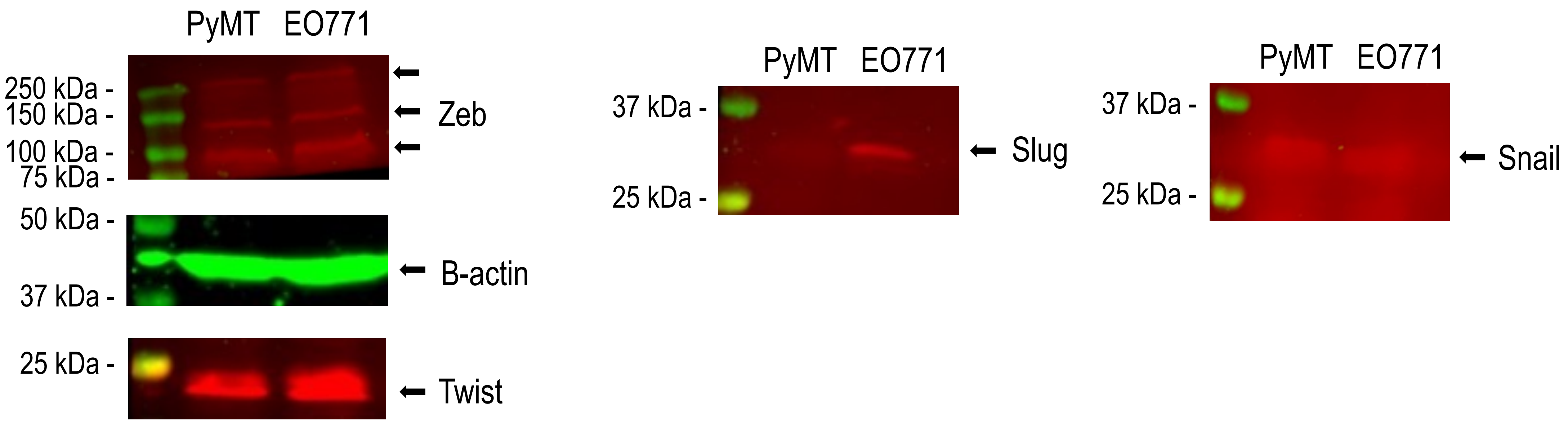

### Fig. S7

Supplementary Figure 7

A

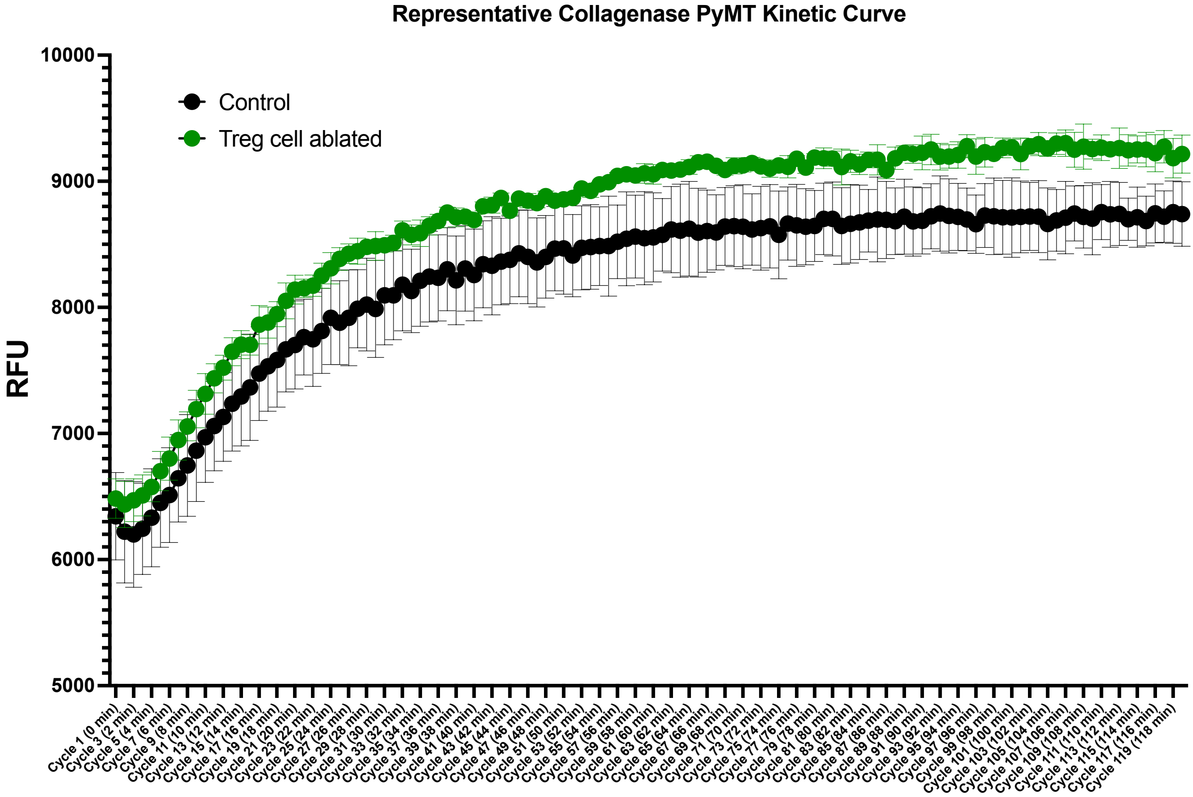


B

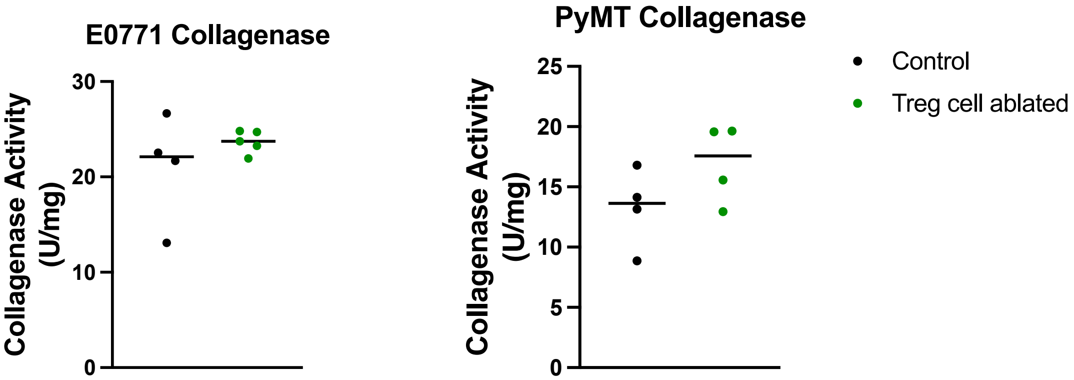


C

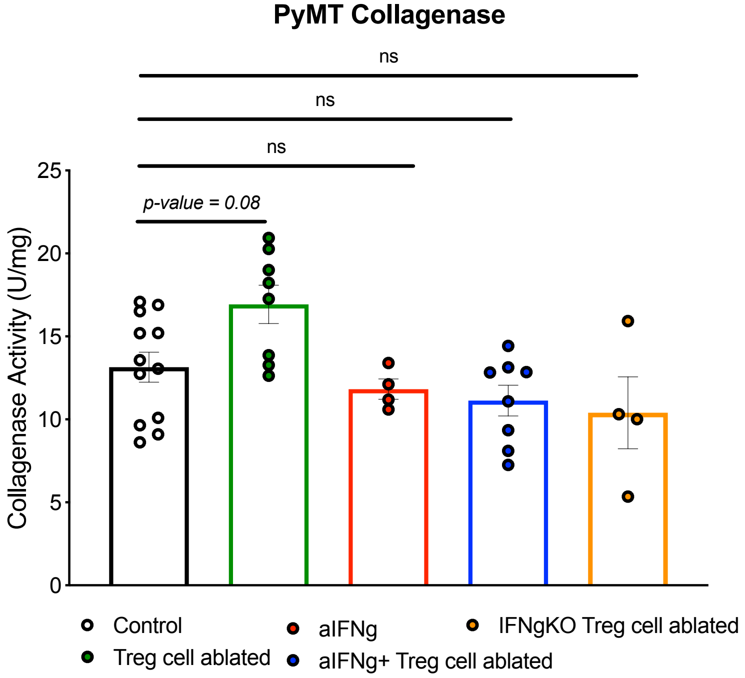

### Fig. S8

Supplementary Figure 8

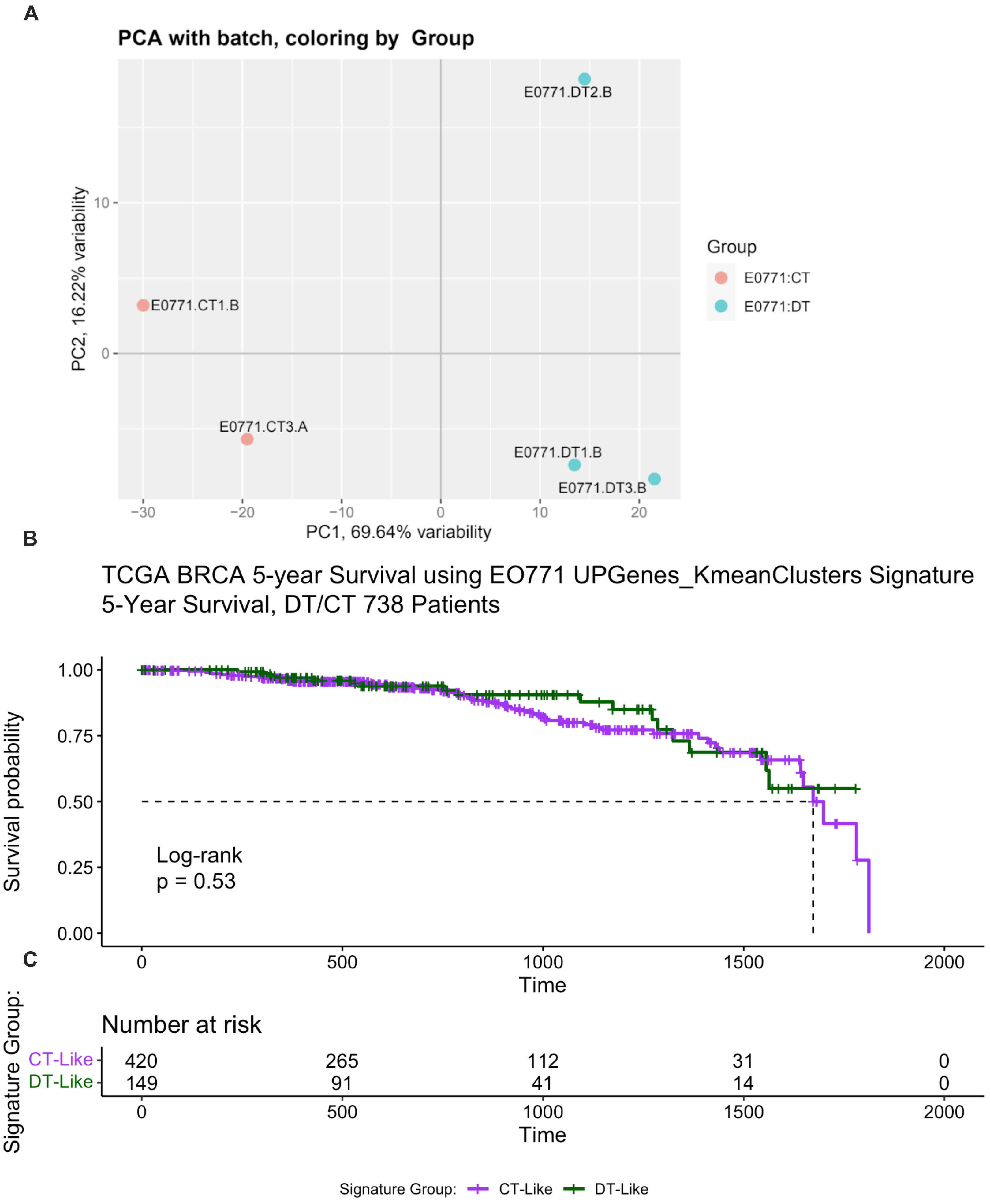

### Fig. S9

Supplementary Figure 9

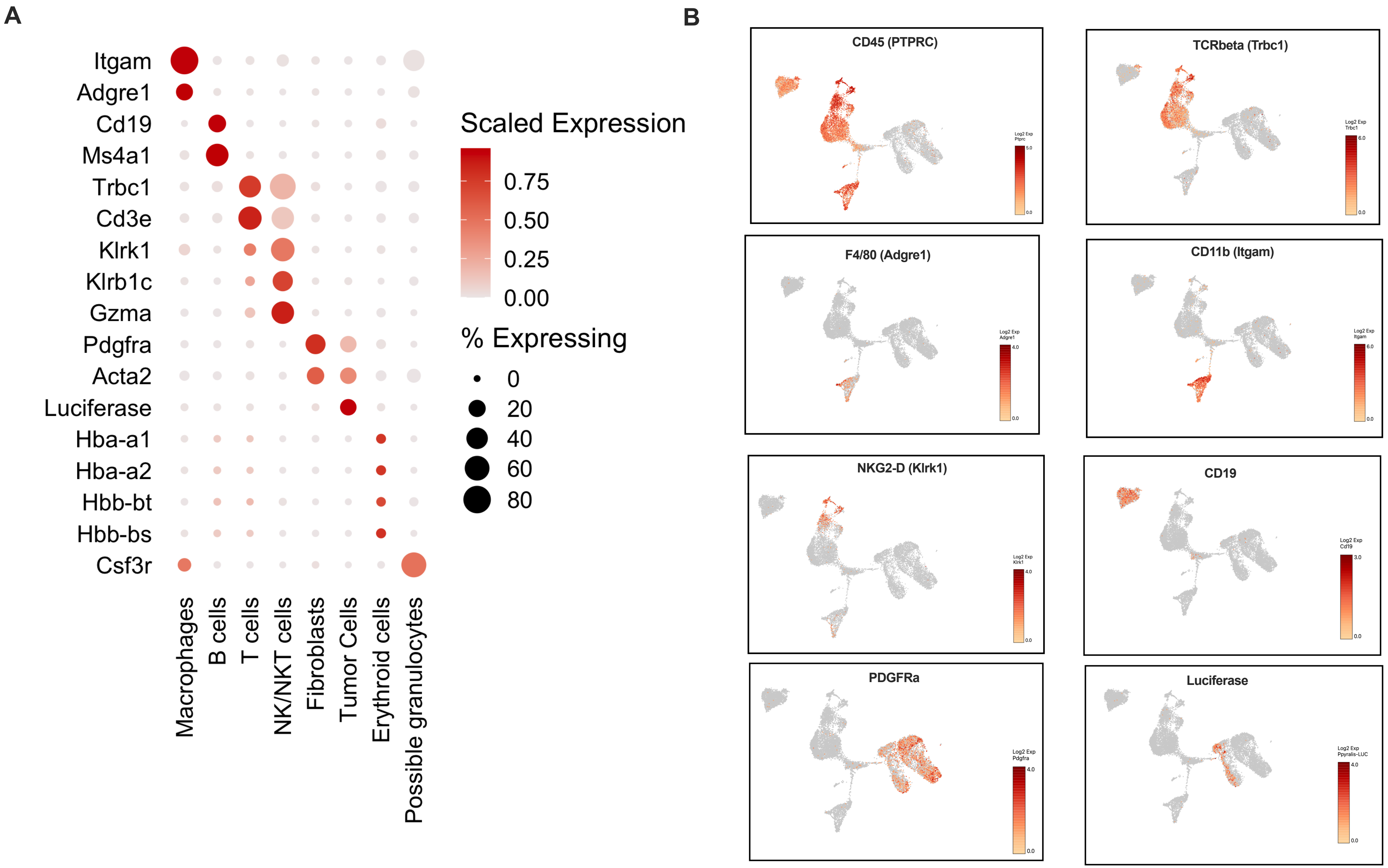
