## Supplementary material for "Tumor cell dissemination is facilitated through regulatory T cell-driven extracellular matrix remodeling": Fig. S6

Supplementary Figure 6

**A** **Unstained Tube**

Tumor Naive Whole Blood

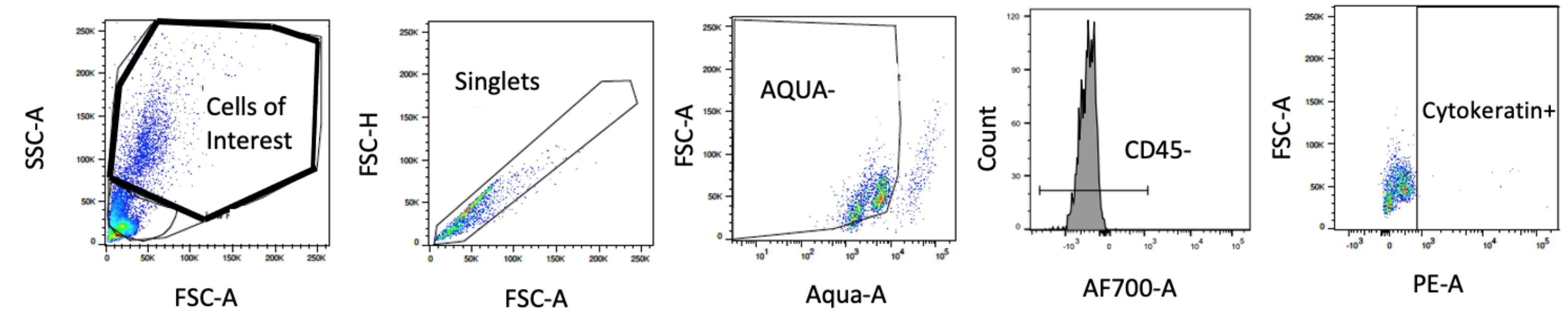


**B** **Stained Tube**

Tumor Naive Whole Blood

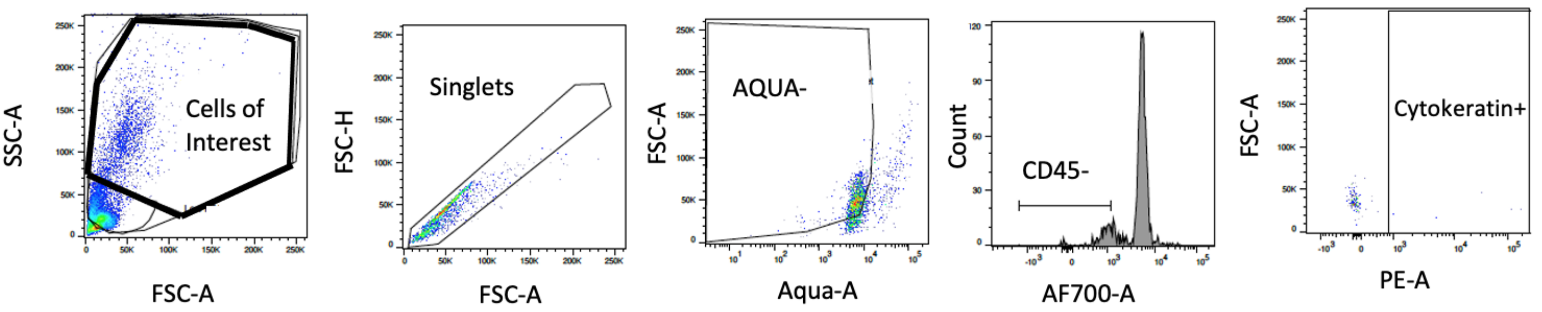


**C** **Stained Tube**

Tumor Naive Whole Blood + Tumor Cell Spike In

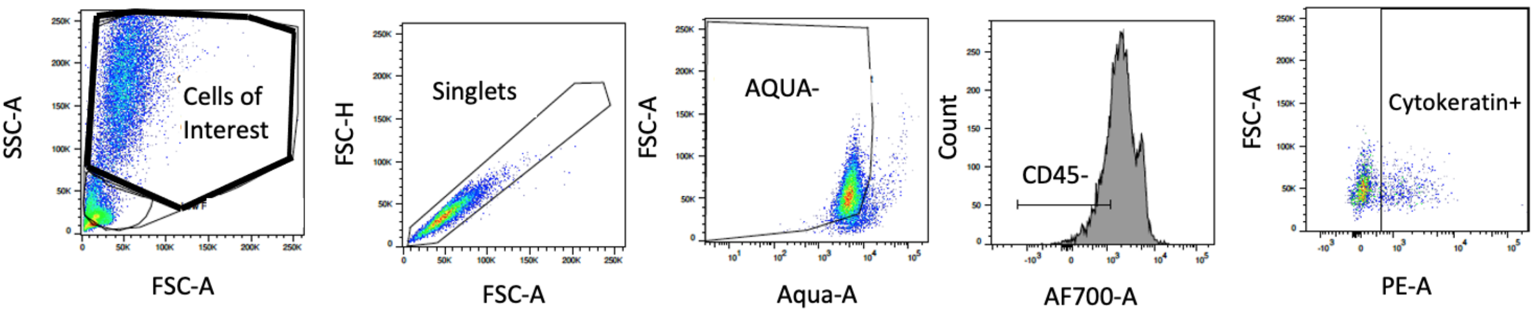
